# A therapeutic small molecule lead enhances γ-oscillations and improves cognition/memory in Alzheimer’s disease model mice

**DOI:** 10.1101/2023.12.04.569994

**Authors:** Xiaofei Wei, Jesus J. Campagna, Barbara Jagodzinska, Dongwook Wi, Whitaker Cohn, Jessica Lee, Chunni Zhu, Christine S. Huang, László Molnár, Carolyn R. Houser, Varghese John, Istvan Mody

## Abstract

Brain rhythms provide the timing and concurrence of brain activity required for linking together neuronal ensembles engaged in specific tasks. In particular, the γ-oscillations (30-120 Hz) orchestrate neuronal circuits underlying cognitive processes and working memory. These oscillations are reduced in numerous neurological and psychiatric disorders, including early cognitive decline in Alzheimer’s disease (AD). Here we report on a potent brain permeable small molecule, DDL-920 that increases γ-oscillations and improves cognition/memory in a mouse model of AD, thus showing promise as a new class of therapeutics for AD. As a first in CNS pharmacotherapy, our lead candidate acts as a potent, efficacious, and selective negative allosteric modulator (NAM) of the γ-aminobutyric acid type A receptors (GABA_A_Rs) assembled from α1β2δ subunits. We identified these receptors through anatomical and pharmacological means to mediate the tonic inhibition of parvalbumin (PV) expressing interneurons (PV+INs) critically involved in the generation of γ-oscillations. Our approach is unique as it is meant to enhance cognitive performance and working memory in a state-dependent manner by engaging and amplifying the brain’s endogenous γ-oscillations through enhancing the function of PV+INs.

## Introduction

The fundamental pathology of Alzheimer’s disease (AD) is largely understood^1^, yet there are no effective drugs to restore cognition and memory in AD patients. Aducanumab (Aduhelm^®^) and lecanemab (Leqembi^®^) were recently approved by the FDA for treatment of AD, however, the approvals were based on significant decreases in the brain amyloid-β (Aβ) pathology, while their ability to reverse cognitive impairment was modest, and side effects still remain a concern.^2^ Cognitive impairment in AD refers to the states of confusion or memory loss that increase yearly in frequency and/or progress in severity. Mild cognitive impairment (MCI) is a condition with more memory problems than what is considered normal for a given age, with symptoms less severe than in AD. About 80% of individuals who fit the definition of amnestic MCI go on to develop AD within 7 years. In a large cohort, pre-diagnostic cognitive impairment and decline with time were present in patients diagnosed 9 years later with AD.^3^ Although age is the primary risk factor for cognitive impairment, other common neurological and psychiatric conditions such as epilepsy^4^, Parkinson’s disease^5^, traumatic brain injury^6^, autism spectrum disorder^7^, depression^8^, schizophrenia^9^, and numerous other morbidities are plagued by loss of cognitive performance. A vast literature has demonstrated that the γ-oscillations generated by fast spiking, parvalbumin-containing interneurons (PV+INs) constitute a fundamental binding scheme and timing mechanism for bringing together brain networks critical for cognitive performance and short-term or working memory.^10–20^ Considerable evidence exists that the neurological and psychiatric disorders mentioned above are also characterized by reduced γ-oscillations.^20,21^

High temporal resolution recording methods, such as magnetoencephalography (MEG)^22^, have made it clear that AD patients starting as early as at the MCI stage have diminished γ-oscillations even before the Aβ load takes full effect.^23,24^ Moreover, it has been well established that almost all AD mouse models have diminished γ-oscillations.^23^ Restoring γ-oscillations in AD patients has yet to be attempted, but remarkably, optogenetic stimulation of PV+INs at a typical γ-oscillation frequency (40 Hz) reduced plaque burden in a mouse model of AD^25^, and sensory (visual and auditory) stimuli ^26,27^ delivered on a daily basis were also effective. These latter methods may well be easily applicable to humans ^28^ for cognitive enhancement therapies. Accordingly, scores of clinical trials are under way^29^ using transcranial magnetic stimulation (TMS), transcranial alternating or direct current stimulation (tACS or tDCS), GammaSense stimulation, transcranial electromagnetic treatment, and intranasal delivery of near infrared light via diodes, all aimed at reducing hyperexcitability and/or improving cognition in AD. However, the timing of the external stimuli will rarely, if ever, coincide with the demand for endogenous γ-oscillations in the brain ^30–32^. A recent study ^32^ failed to replicate the plaque clearing effects of 40 Hz stimulation ^25^ and pointed out the inability of external periodic stimuli to engage γ-oscillations in the brain. We therefore decided to pursue a pharmacological approach to enhance the endogenous γ-oscillations of the brain by reducing the tonic inhibition of PV+INs. This approach should work when these natural rhythms are engaged by the brain’s own circuitry during cognitive demand and working memory.^33–36^

## Results

### The specific GABA_A_R partnership responsible for the tonic inhibition of PV+INs

In the Cys-loop superfamily of ligand gated ion channels, the GABA_A_Rs stand out as the most diverse based on subunit composition and pharmacology. From nearly 500,000 possible heteropentameric combinations of 19 different subunits, due to specific receptor partnership rules, there are only about 26 naturally occurring subtypes in the brain^37^, but these may be augmented by the recently discovered diversity of receptor assembly.^38^ As various subunits impart specific biophysical properties such as channel open times, desensitization, deactivation, GABA affinity and potency, they also define a wide array of pharmacological properties that modulate the function of these receptors.^37^ The following native δ-GABA_A_Rs subunit combinations are present in the brain:^39^ α6β2δ are abundant on cerebellar granule cells; α4β2/3δ are most prominent in the forebrain where they are found on dentate gyrus granule cells, various thalamic neurons, medium spiny striatal neurons, neocortical pyramidal cells and other brain regions; and presumably α1-subunit-containing receptors are found on δ-GABA_A_Rs PV+INs^40–44^ and possibly on cortical and hippocampal INs termed neurogliaform cells^41,43,45^, which actually decouple pyramidal cells from γ-oscillations.^46^ An intriguing finding regarding the brain’s native δ-GABA_A_Rs is the assembly of δ subunits only with three different α subunits (α1, α4, and α6). When the α4 or α6 subunits are genetically deleted from mice, the δ subunits fail to pair up with existing α1 subunits, even though the cells with missing α4 or α6 subunits possess plentiful α1 subunits. Therefore, cells with deleted α4 or α6 subunits also become δ subunit knockouts.^43,47^

Using conventional microscopy we have previously shown the colocalization of δ-subunits in PV+INs of the CA3 region^42^. We have now extended these findings using confocal microscopy to the localization of α1 subunits in the PV+INs of this region (Figure 1a). We also confirmed the colocalization of α1 and δ subunits in the characteristically pyramidal shaped PV+INs (basket cells) of the dentate gyrus (Supplementary Figure S2a). We then set out to examine the strength of α1/δ subunit partnership in the same neurons when δ-subunits were absent following genetic deletion. We observed strong membrane staining of α1 subunits in PV+INs of WT mice over the entire extent of the cells (Figure 1b), consistent with an extrasynaptic localization of these receptors. In sharp contrast, α1 subunit surface labeling was decreased and the α1 subunits appeared to be trapped in the cytoplasm in homozygous δ-subunit knock-out (Gabrd-/-) mice (Figure 1c), indicating that the α1/δ subunit partnership is a key element for ensuring surface expression of these subunit combinations. We have also identified the type of β subunit present in the PV+IN GABA_A_R assembly as the β2 variant (Supplementary Figure S2b), while finding no measurable staining for β3 subunits (Supplementary Figure S2c). These data strongly corroborate the subunit composition of the extrasynaptic GABA_A_Rs of PV+INs, most likely to be responsible for their tonic GABA conductance, as having the α1β2δ subunit composition, consistent with a previous report.^44^ At this time, we cannot ascertain the stoichiometry of the extrasynaptic receptors of PV+INs, as they may show similarities with the recently identified unusual stoichiometries of extrasynaptic receptors with α4β3δ subunit composition.^38^

**Figure 1.**
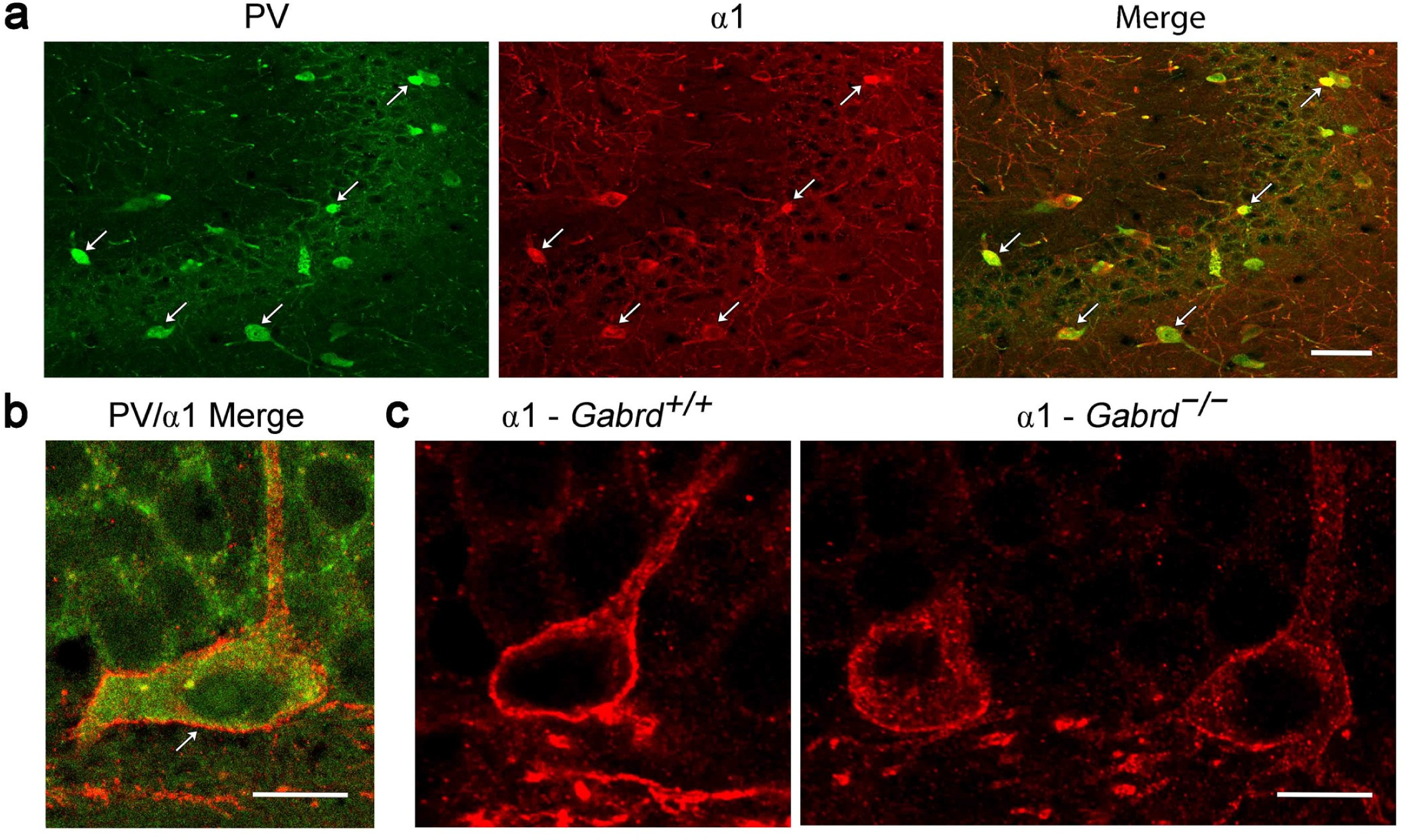
*α*1 subunit is prominently expressed in PV+ interneurons in CA3 and in similar interneurons along the base of the dentate granule cell layer. **a.** Many PV+INs in CA3 express the α1 subunit (examples at arrows). Their distribution closely resembles that of δ subunit-expressing PV+INs in CA3 (described previously^42^). **b.** PV+INs along the base of the granule cell layer also exhibit strong α1 subunit labeling along their cell surface (arrow). **c.** In *Gabrd^+/+^* animals, α1 subunit labeling is similarly expressed on the surface of these interneurons, with lighter, diffuse labeling within the cytoplasm. In contrast, in *Gabrd^-/-^* animals, many α1-labeled interneurons have reduced surface labeling and exhibit increased punctate labeling within the cytoplasm. This suggests that loss of the δ subunit leads to altered localization of the α1 subunit in these interneurons, consistent with a favored partnership of the δ and α1 subunits. Scale bars: **a** = 50 µm; **b,c** = 10 µm.

### Pharmacological specificity and selectivity of the α1β2δ GABA_A_Rs underlying tonic inhibition of PV+INs

There are notable differences between the properties of δ-GABA_A_Rs depending on which α subunits are present in the receptor. In expression systems (*Xenopus* oocytes, HEK cells) the potency of GABA is higher at α4 versus α1 containing δ-GABA_A_Rs. The EC50 of GABA at α4δ-GABA_A_Rs is in the high nM range,^48–50^ while at α1δ-GABA_A_Rs it is about 10-fold higher, in the low μM range.^48–53^ This raises an interesting question about when exactly the α1δ-GABA_A_Rs of PV+INs^40–44^ and those on cortical and hippocampal neurogliaform cells^41,43,45^ become activated? Apparently, the high activity of PV+INs during γ-oscillations,^10,11,17,54^ generates sufficient (low μM) levels of GABA for the α1δ-GABA_A_Rs of PV+INs to become activated, thus controlling γ-oscillation frequency and amplitude. However, during γ-oscillations the same receptors on neurogliaform cells will not be active, as these cells do not participate in γ-oscillations.^46,55,56^ There are also differences in the effects of positive allosteric modulators (PAM) on δ-GABA_A_Rs that are determined by the presence of different α subunits. Whereas the effects of neurosteroids such as allopregnanolone and THDOC appear to be similar irrespective of the α subunits, the effects of ethanol and the δ-GABA_A_R selective modulator DS2 show different activities in expression systems compared to the native receptors in the brain. In expression systems DS2 is equally potent at α4- and α1δ-GABA_A_Rs ^49,51,57,58^, yet DS2, in sharp contrast to allopregnanolone, does not alter γ-oscillations in slices with native α1δ-GABA_A_Rs ^40,42^. It is interesting to note that in oocytes, tracazolate, a nonbenzodiazepine anxiolytic, considerably increases both the potency and efficacy of GABA δ-GABA_A_Rs, particularly in receptors with α1β2δ composition ^52^. As no studies were published on the potential effects of tracazolate on native neuronal GABA_A_R assemblies, we decided to test its effects on the tonic and phasic inhibition in PV+INs to ascertain also by pharmacological means whether GABA_A_Rs α1β2δ composition were present in these neurons. As shown in Figure 2a, tracazolate at a concentration of 10 μM, which was found to be the effective concentration in oocytes ^52^, also enhanced the tonic currents mediated by α1β2δ GABA_A_Rs recorded in PV+INs. The summary data (Figure 2a) indicate that the effect was significant (Control: 0.86 ± 0.45 A/F vs 10 μM tracazolate: 1.56 ± 0.36 A/F, mean ± SD, n=5 cells, n=5 mice; an 82% increase, p=0.0139, paired t-test).

**Figure 2.**
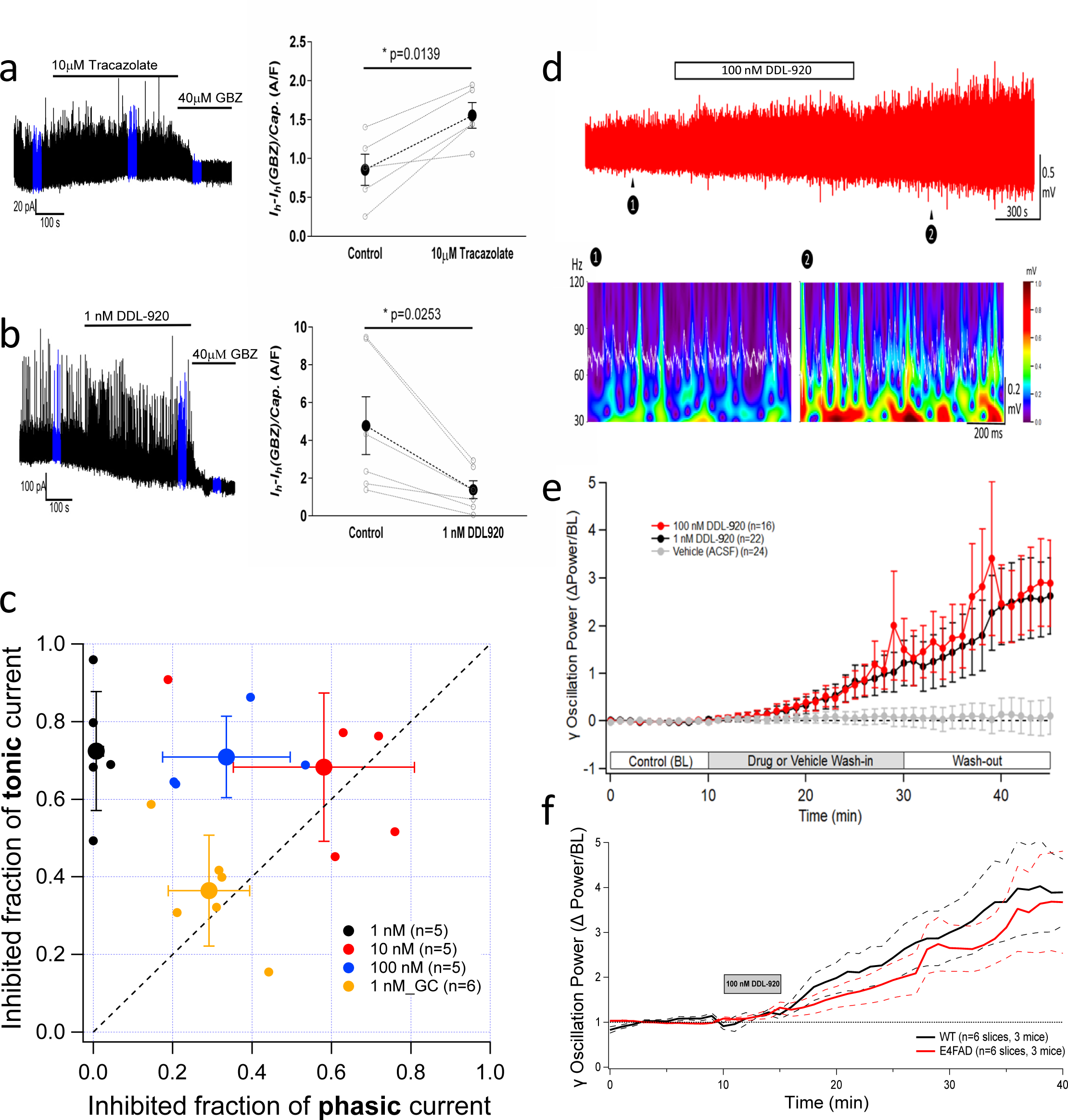
Pharmacological characterization of DDL-920 in PV+INs and its effects on γ-oscillations *in vitro*. **a**, The effects of tracazolate, a compound known to potentiate GABA_A_Rs composed of α1β2δ subunits in Xenopus oocytes, on the tonic GABA_A_R-mediated currents of identified PV+INs. *Left panel*: Raw traces of a voltage-clamp recording at Vh=0 mV of the tonic GABA_A_R-mediated currents before, during and after the perfusion of 10 μM tracazolate (*horizontal bar*). The perfusion of 40 μM gabazine (*GBZ*) was used to block all GABA_A_Rs. The *blue* segments in the recording indicate the 30 s epochs used for the tonic and phasic current analysis. *Right panel*: Summary data from 5 recordings where the tonic GABA_A_R-mediated currents normalized to the capacitance of the cells are plotted before (*Control*) and during (*10 μM tracazolate*) the perfusion of the drug. The individual experimental results are illustrated with light grey open circles connected by dotted lines, while the averages (± SEM) are indicated by solid black circles connected by a black dotted line. The p-value indicates a significant increase in the tonic current as determined using the paired Wilcoxon signed rank test. **b**, The effects of DDL-920 on the tonic GABA_A_R-mediated currents of identified PV+INs. *Left panel*: Raw traces of a voltage-clamp recording at Vh=0 mV of the tonic GABA_A_R-mediated currents before, during and after the perfusion of 1 nM DDL-920 (*horizontal bar*). The perfusion of 40 μM gabazine (*GBZ*) was used to block all GABA_A_Rs. The *blue* segments in the recording indicate the 30 s epochs used for the tonic and phasic current analysis. *Right panel*: Summary data from 6 recordings where the tonic GABA_A_R-mediated currents normalized to the capacitance of the cells are plotted before (*Control*) and during (*1 nM DDL-920*) the perfusion of the drug. The individual experimental results are illustrated with light grey open circles connected by dotted lines, while the averages (± SEM) are indicated by solid black circles connected by a black dotted line. The p-value indicates a significant reduction of the tonic current as determined using the paired Wilcoxon signed rank test. **c,** Plot of the inhibited fraction (efficacy of inhibition, see Methods for details) of the tonic current vs the same for the phasic current by DDL-920 in cortical PV+INs. The small colored dots are from individual cell recordings while the large dots are the average values with the SD bars extending in both directions. The concentrations of the compounds are color-coded as per the legend. The dashed line has a unity slope and thus indicates no specificity of the compound for either tonic or phasic inhibition. (*orange* data points are from dentate gyrus granule cells (GC) obtained at 1 nM DDL-920). **d,** Effects of 100 nM DDL-920 perfused for 15 min on *in vitro* γ-oscillations in the hippocampal CA3 pyramidal layer. The Morlet wavelets (*bottom panel*) show the large enhancement of the magnitude of 40-50 Hz oscillations during 1 s epochs before (❶) and after (❷) the application of DDL-920. **e,** Comparative effects of 20 min perfusions of 1 or 100 nM DDL-920, or vehicle (aCSF) on the power (RMS averaged during 60 s epochs) of *in vitro* γ-oscillations (n’s represent the number of slices). The changes are expressed relative to the power measured during the baseline recording period (10 min). **f,** Effects of 5 min perfusions of 100 nM DDL-920 on slices obtained from WT and AD model mice. Measurements as in e. The solid lines represent the averages, the dashed lines are the SD envelopes of the averages. There are no significant differences between the enhancement of γ-oscillation power by DDL-920 in the two genotypes.

The pharmacological effect of tracazolate, a positive allosteric modulator (PAM) of α1β2δ GABA_A_Rs expressed in oocytes, was consistent with the same combination of GABA_A_R subunits mediating the tonic GABAergic inhibition in murine cortical and hippocampal PV+INs. Thus, we next wanted to test the effects of our potential negative allosteric modulator (NAM) compound DDL-920 on the same neurons. The structure and synthesis of DDL-920 are shown in our published international patent application^59^. At concentrations as low as 1 nM, DDL-920 reduced the large tonic currents elicited by 5 μM GABA in PV+INs (Figure 2b; Control: 4.78 ± 3.75 A/F vs 1 nM DDL-920: 1.39 ± 1.16 A/F, mean ± SD, n=6 cells, n=6 mice; a 71% decrease, p=0.0253, paired t-test). As tonic inhibition in PV+INs is mediated by extrasynaptic α1β2δ GABA_A_R (see previous section), while phasic inhibition is generated by fast synaptic α1β2γ2 receptors^60^, a differential effect of DDL-920 on the tonic vs phasic currents will indicate how specific this compound is for the native α1β2δ GABA_A_Rs of PV+INs. In a different set of experiments, we measured the compound’s tonic vs phasic specificity by plotting the fraction blocked of the tonic current vs that of the phasic current (Figure 2c). The fractions blocked are calculated as: 1 - (current in drug/current in control). The specificity for the inhibition of the tonic vs phasic current was calculated as: fraction blocked (tonic/phasic). For 1 nM DDL-920 this value is: 0.792/0.011, i.e., 72-fold. We further examined the selectivity of DDL-920 for the α1β2δ GABA_A_Rs found in PV+INs compared to the α4β2δ GABA_A_Rs mediating the tonic inhibition in dentate gyrus granule cells^61^. DDL-920 at 1 nM was approximately 2-fold more selective for the tonic currents mediated by α1β2δ GABA_A_Rs in PV+INs than those of dentate gyrus granule cells mediated by α4β2δ GABA_A_Rs (Figure 2c). We did not perform a full dose-response analysis of the effects of DDL-920 on the tonic GABA currents of PV+INs, as we wanted to stay in the range of the concentrations we found in the brains of mice after subcutaneous (SQ) or oral administration at a standard dose of 10 mg/kg (Supplementary Figure S3a&b). However, from the three doses of DDL-920 examined on the tonic inhibition of PV+INs in vitro (Figure 2c), it appears that the compound already reaches full efficacy at 1 nM.

We have also engaged in some molecular modeling of DDL-920 binding to the available GABA_A_R homo- and heteropentameric structures resolved through protein crystallography or cryo-EM. The docking of the DDL-920 molecule to the homomeric β3 GABA_A_R (pdb: 4COF) is shown on Supplementary Figure S4a. The binding gives a good lead finder (LF) rank score of - 10.34 which is further improved with the addition of the GABA_A_R δ subunit (pdb: 7QND; Supplementary Figure S4b). Unfortunately, the precise structural model for the unique α1β2δ GABA_A_R combination found on PV+INs is not available, but our docking modeling shows that the introduction of the δ subunit provides additional binding sites for DDL-920. The use of the β3 GABA_A_R subunit in our model instead of the β2 is not a great concern, as in their transmembrane regions where DDL-920 appears to bind, there is a nearly 100% homology between the two subunits.

### Effects of DDL-920 on γ-oscillations in vitro

In horizontally cut hippocampal slices γ-oscillations can be readily recorded just below the CA3 pyramidal cell layer in the presence of 50 nM kainic acid^40,42,62,63^ (Figure 2d). These oscillations recorded *in vitro* correspond well to those recorded *in vivo* during spatial reference memory.^62^ In this region, the only cells with δ-GABA_A_Rs are the PV+INs^40^. Our previous studies have produced multiple lines of evidence substantiating the expression levels of δ-GAB_A_R receptors on PV+INs effectively controlling γ-oscillation amplitude and frequency recorded *in vitro*. (1) Genetic deletion of δ-subunits and the consequent loss of tonic inhibition onto these INs in Gabrd-/- mice has major potentiating effects on the frequency of *in vitro* CA3 γ oscillations ^63^.

More direct evidence came from our studies using conditional knockouts to specifically delete or reduce δ-GABA_A_R expression *only* in PV+INs^42,64^ resulting in an increased frequency and power of γ-oscillations. Already a 30% reduction in δ-GABA_A_R in PV+INs significantly increases γ-oscillation frequency.^42^ Conversely, potentiation of the δ-GABA_A_R-mediated tonic conductance with neurosteroids lowered the power of γ-oscillations.^40^ (2) During pregnancy, δ-GABA_A_R levels on PV+INs decline by about 60% without any change in their PV content. Concomitantly, the frequency of γ-oscillations measured in *ex vivo* slices from pregnant mice increases considerably.^40^ *In vivo*, these altered γ-oscillations are counterbalanced by the increased levels of allopregnanolone during pregnancy to potentiate the function of the reduced number of δ-GABA_A_Rs on PV+INs just sufficiently to restore the normal (lower) range of γ-oscillations frequency.^40^

Based on these findings, our next step was to examine the effects of DDL-920, the NAM of the α1β2δ-GABA_A_Rs of PV+INs, on γ-oscillations recorded *in vitro* (Figure 2d). The power of the γ-oscillations was evaluated as the root mean square (square root of the mean squared, RMS) of the 30-120 Hz filtered local field potential (LFP). This value was computed for every 60 s of the recordings (Figures 2e,f). We thus demonstrated similar increases in γ oscillatory power following perfusion of 1 or 100 nM DDL-920 for 20 min (Figure 2e), while perfusion of the artificial cerebro-spinal fluid (ACSF) with no drugs had no effects on the power of γ-oscillations (Figure 2e). The change in power over baseline was calculated as the RMS during each of 60 s – the mean RMS during 10 min baseline divided by the mean RMS during the 10 min baseline, as in Figure 2e. Thus, a value of 1 corresponds to a 100% increase in γ-oscillatory power over that recorded during the 10 min baseline. After 15 min of washout, the power of the γ-oscillations was still increased by ∼2.5-fold after administration of 1 or 100 nM DDL-920 for 20 min (Figure 2e). This finding indicates that, at least *in vitro*, the maximal efficacy of DDL-920 on increasing the power of the γ-oscillations is already reached at 1 nM, a concentration that blocks >75% of tonic inhibition of PV+INs with only a negligible effect on their phasic inhibition (Figure 2c).

Once we demonstrated the boosting effects of DDL-920 on γ-oscillations in *ex vivo* slices of WT mice, we went on to examine its effects on oscillations recorded in slices from our AD model mice (see Methods for details). As shown in Figure 2f, we found similar large enhancements in the power (RMS) of γ-oscillations recorded in the hippocampal CA3 pyramidal layer of slices after a 5 min perfusion of 100 nM DDL-920 of WT (n=6 slices, 3 mice) and AD model mice (n=6 slices, 3 mice). These promising results in AD mice lead us to test the effects of DDL-920 on γ-oscillations recorded *in vivo*.

### Effects of DDL-920 on γ-oscillations in vivo

Based on our previously published findings, we had good reason to believe that reducing the effectiveness of tonic inhibition of PV+INs *in vivo* will boost the power of γ-oscillations. We noticed that the expression levels of α1δ-GABA_A_Rs of PV+INs fluctuate during the ovarian cycle and the γ-oscillation frequency and power fluctuate correspondingly.^64^ For example, during the estrus phase of the cycle, when the δ-GABA_A_R expression in PV+INs is diminished by about 30%, the amplitude of γ-oscillations is increased by a commensurate amount.^64^ Notably, similar alterations in γ-oscillations are also found in healthy women during the menstrual cycle.^65^.

The inverse correlation between δ-GABA_A_R expression in PV+INs and γ-oscillation power *in vivo* is promising, but prior to examining the potential effectiveness of DDL-920 on γ-oscillations *in vivo*, we had to ensure that the drug was permeable through the BBB, and that it had a reasonable pharmacokinetic (PK) profile following subcutaneous (SQ) or oral (PO) administration. Mice (n=4/dose) were given DDL-920 SQ at 1, 5, 10 mg/kg and PO at 10 mg/kg. Plasma and brain were collected 1 and 3 hrs after drug administration. Targeted liquid LC-MS/MS assay was developed for measurement of DDL-920 concentrations. Supplementary Figure S3a shows that 1 hour after 10 mg/kg DDL-920 SQ injection the maximal concentration (C_max_) of the drug in the brain reaches about 25 nM, but it declines to ∼10 nM after 3 hours. The same dose 1 hour after oral gavage reaches a C_max_ of ∼30 nM but lasts for a longer period (Supplementary Figure S3b), and after oral pipette administration reaches a C_max_ of ∼23 nM. Furthermore our *in vitro* ADME data show that DDL-920 has good kinetic solubility at >100 μM in water, plasma stability with t_1/2_ > 180 min with a free unbound fraction (F_u_) of 34% in brain tissue homogenates (see Methods, and Supplementary Table S1), suggesting that the concentration of unbound DDL-920 in the brain is above the *in vitro* efficacious dose for γ-oscillation enhancement seen in brain slices (Figure 2e). These findings made us confident that a few hours following SQ or oral administration, DDL-920 will reach the brain in sufficient amounts to antagonize tonic inhibition of PV+INs (Figure 2b&c) and thus enhance γ-oscillations.

We first administered DDL-920 (n=4) or saline (n=3) to adult WT mice in which bilateral hippocampal electrodes were implanted in the CA1 region to record LFP 24/7 using a chronic tethered recording setup.^64,66^ We injected 10 mg/kg SQ during the middle of their light cycle (at around 12 PM) and monitored the hippocampal LFP for several hours prior and after the injection. The 180 s segments used for the analyses were randomly selected based on consistent amplitude θ-oscillations from periods of 3 hrs before and 3 hrs after injections, all chosen from segments where there was above average power (RMS) in the θ frequency (5-12 Hz). We chose the θ frequency to guide us for the γ-oscillation segments because of the tight phase amplitude coupling (PAC) between the phase of θ-oscillations and γ-oscillation amplitude, a process considered critical for cognition and short-term/working memory.^20,67–69^ In Figures 3a-g we illustrate several properties of the γ-oscillations and their PAC to θ-oscillations in an animal that received two SQ DDL-920 injections 3 weeks apart and a SQ saline injection 3 days after the first DDL-920 administration. Figure 3a shows 1 s long traces to illustrate typical examples of the θ- and γ-oscillations recorded in this mouse during the three pre-injection periods while Figure 3b shows similar traces after the DDL-920 injections. The times before and after the injections are indicated in the legend. The modulation index of the γ-oscillation amplitude by the phase of the θ-oscillations at a given frequency^68,69^ calculated for the entire duration of the 180 s recordings shows a remarkable increase following the two administrations of DDL-920 and no change after saline administration. We also examined the phase of the θ-cycle during which the θ-oscillations have the largest amplitude. When measured in 10 Hz bins, the highest amplitude θ-oscillations always corresponded with the peak (0 radians on a scale of −π to +π) of the θ-cycle regardless of θ-oscillation frequency (Figure 3d). This finding is noteworthy, as it means that in spite of the considerable DDL-920-induced increase in the amplitude of γ-oscillations (Figures 3b,f,g), their PAC to θ-oscillations remained constant. We have also plotted the instantaneous frequencies (Figure 3e) and amplitudes (Figure 3f) of the θ- and γ-oscillations recorded during the 180 s representative epochs. The frequency distributions were remarkably unaltered (Figure 3e) showing peaks of 8-9 Hz for θ-oscillations and two peaks (∼40 Hz and ∼80 Hz) for the low and high γ-oscillations, respectively. Accordingly, while DDL-920 increased the power of γ-oscillations, it did not alter their frequencies. Figure 3f shows the log-scaled peak-to-peak amplitudes of the θ- and γ-oscillations plotted against their instantaneous frequencies. The amplitudes of the θ-oscillations increased after each of the 3 injections, including after saline. In contrast, γ-oscillation amplitudes increased (by 15-55%) over their entire frequency range following DDL-920 injections and decreased after saline injection. We also plotted the histograms of the instantaneous power (RMS) of the θ- and γ-oscillations during the same recording periods (Figure 3g). However, the spectrograms over long time periods (4 hrs before, and 4 hrs after the SQ injections) of the γ-oscillations for the same injections (Supplementary Figures S4-6) better illustrate the comparisons between the power of γ-oscillations during the pre- and post-injection times. Summary data from the 4 WT mice injected with DDL-920 and the 3 mice injected with saline are depicted in Figure 3h. The lines fitted to the data indicate that only the θ-phase γ-amplitude modulation index (5.8-fold) and the γ-oscillation power (RMS; 1.48-fold) were enhanced by DDL-920 when compared to saline injections, whereas the frequencies of θ- and γ-oscillations were unaffected. No other parameter was affected by saline injection, except the θ-γ PAC modulation index (MI) which became reduced by 42% (Figure 3h). This may be due to some mild stress induced by the handling and SQ injections of the WT animals. However, following the SQ DDL-920 or saline injections we did not observe any potential adverse side effects such as hyperexcitability, abnormal spiking, or changes in motor behavior. We have measured the aggregate motor activity patterns derived from the video recordings over 3 hrs before and 3 hrs after the DDL-920 or saline injections. The mean ± SD of RMS (in video pixel units) of the motor activities (n=8 injections, n=4 mice) prior to DDL-920 administration was 6.85 ± 5.56. This was not significantly different than the value measured after the injection (7.12 ± 2.42; p=0.38, Wilcoxon signed rank test). Similarly, the pre- and post-saline injection activity levels (n=6 injections, n=3 mice) did not differ from each other (pre: 8.02 ± 5.72 vs post: 8.55 ± 4.49; p=1, Wilcoxon signed rank test).

**Figure 3.**
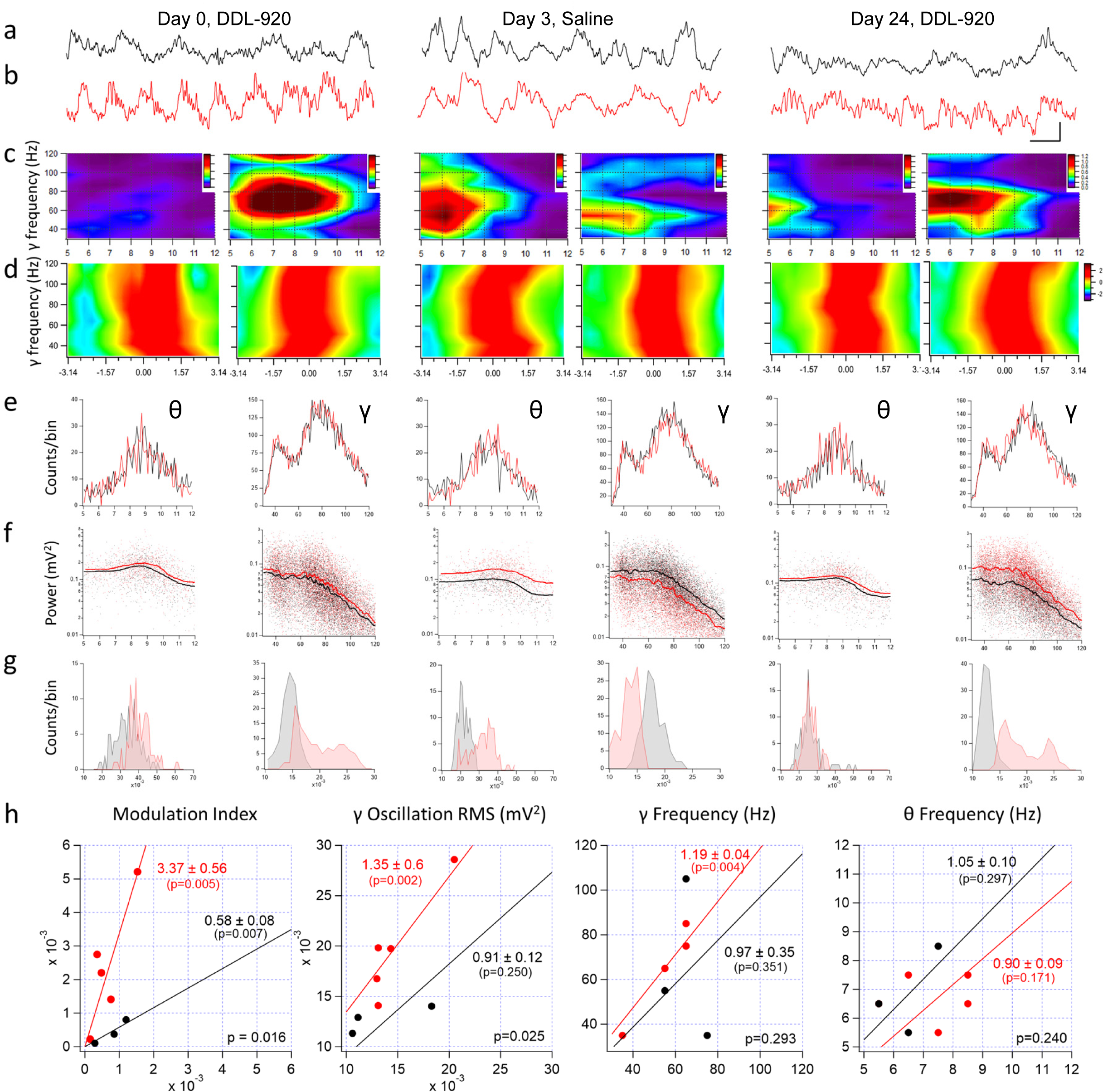
Effects of DDL-920 (10 mg/kg) SQ *in vivo* administrations on hippocampal θ- and γ-oscillations in wild type mice. **a-g:** The effects of two DDL-920 injections >3 weeks apart in a WT mouse (*in red*), with an intercalated saline injection (*in black*) on day 3 after the first DDL-920 injection. **a,** The traces are raw recordings of 1 s long epochs taken between 1-3 hrs prior and **b**, ∼4 hrs, ∼2 hrs, and ∼2.5 hrs, after the DDL-920, saline, and DDL-920 injections, respectively. Calibration bars are 100 ms and 0.2 mV and refer to all traces in a and b. **c-g**: Analyses of θ-γ frequencies, frequency-amplitude (FAC) and phase-amplitude coupling (PAC) in 120 s segments randomly extracted during the indicated periods before and after the injections. **c**, Modulation index (MI) matrix for θ-γ frequencies during the corresponding 120 s before and after injection epochs. The MI shows by how much γ amplitude for each θ frequency bin deviates from a uniform distribution (Kullback-Leibler distance between two distributions^68,69^). Each of the plot pairs are on the same scale. Note the large increases in the MI only after DDL injections. **d**, PAC of θ-γ oscillations with θ phase represented from trough (-π) to peak (0) and back to trough (+π). The values are z-scores calculated within each 10 Hz frequency bin from 30-120 Hz. Despite large increases in γ oscillation amplitudes, their coupling to θ phase remains unaltered after DDL. **e,** Histograms of the instantaneous frequencies (in Hz) of the θ-(*left panels*) and γ-oscillations (*right panels*) before (*black*) and after (*red*) injections showing no differences in the frequencies. Note the presence of both low and high γ-oscillation frequencies. **f**, Instantaneous amplitude vs frequency (in Hz) of θ-(*left panels*) and γ-oscillations (*right panels*) before (*black*) and after (*red*) injections showing increases in the γ-oscillation amplitudes across all frequencies after DDL injections. Please note the log scale of the amplitudes. **g,** Histograms (at bin widths of 0.001 mV^2^) of RMS measured over 120 s of θ-(*left panels*) and γ-oscillations (*right panels*) before (*black*) and after (*red*) DDL injections showing large increases in the γ-oscillation RMS after DDL administrations. **h.** The parameters indicated on the top of the graphs are plotted as post vs pre-injection values (DDL-920: *red* n=5 mice, 5 injections; saline: *black*, n=3 mice, 3 injections). The slopes of the linear regressions through the origin are shown next to the respective regression lines. The p-values in the lower right corners of the plots refer to the comparisons between the two slopes. The slopes (± SD) of the MI and of the γ-oscillation RMS are significantly larger after DDL injection than after saline, indicating that SQ injection of 10 mg/kg DDL-920 potentiated these two measures. In contrast, the frequencies of the θ- and γ-oscillations were not altered by the DDL-920 administration. The p-values shown in the respective colors under the values of the slopes denote the differences between the individual regression line slopes and a slope = 1, i.e., the “no change” line. The frequencies were plotted as the midpoint from the largest values within 10 Hz bins (γ-oscillations) and 1 Hz bins (θ-oscillations), and therefore, some points overlap in the plots.

### Effects of DDL-920 on AD model mice cognitive/memory performance

Encouraged by the superior PK of DDL-920 after oral administration (Supplementary Figure S3b, Supplementary Table S1) we decided to examine its effects in AD model mice following oral administration with a pipettor. Supplementary Figure S8a&b shows that this route of administration increases γ-oscillation power in AD model mice for about 4.5 hrs, indicating a coarse measure of its pharmacodynamic effect. We administered DDL-920 by this method to 3 AD mice implanted with bilateral recording electrodes in the CA1 region of the hippocampus.

The recordings were chosen just as for the WT mice (see above) and 1 s long epochs both before and after the DDL-920 (10 mg/kg) administration to one of these mice is shown in Figure 4a. The instantaneous amplitudes of the θ-oscillations were unaffected, but those of γ-oscillations were increased (15-35%) over all frequencies, with a significant shift to higher values of the calculated γ/θ amplitude ratios (Figure 4b). In the same mouse, an oral saline administration did not produce changes in any of these three parameters (Figure 4c). Similar results were obtained in the other 2 AD mice. Unfortunately, we could not obtain the PAC parameters in these mice, as establishing the location of the electrodes (i.e., ascertain the troughs and peaks of the θ-oscillations) relies on recording sharp wave ripples.^66^ These events were only a few in the AD mice, as they appear to have been replaced by larger amplitude spiking activity. The characterization of this phenomenon was beyond the scope of our investigation as similar findings have recently been reported.^70^

**Figure 4.**
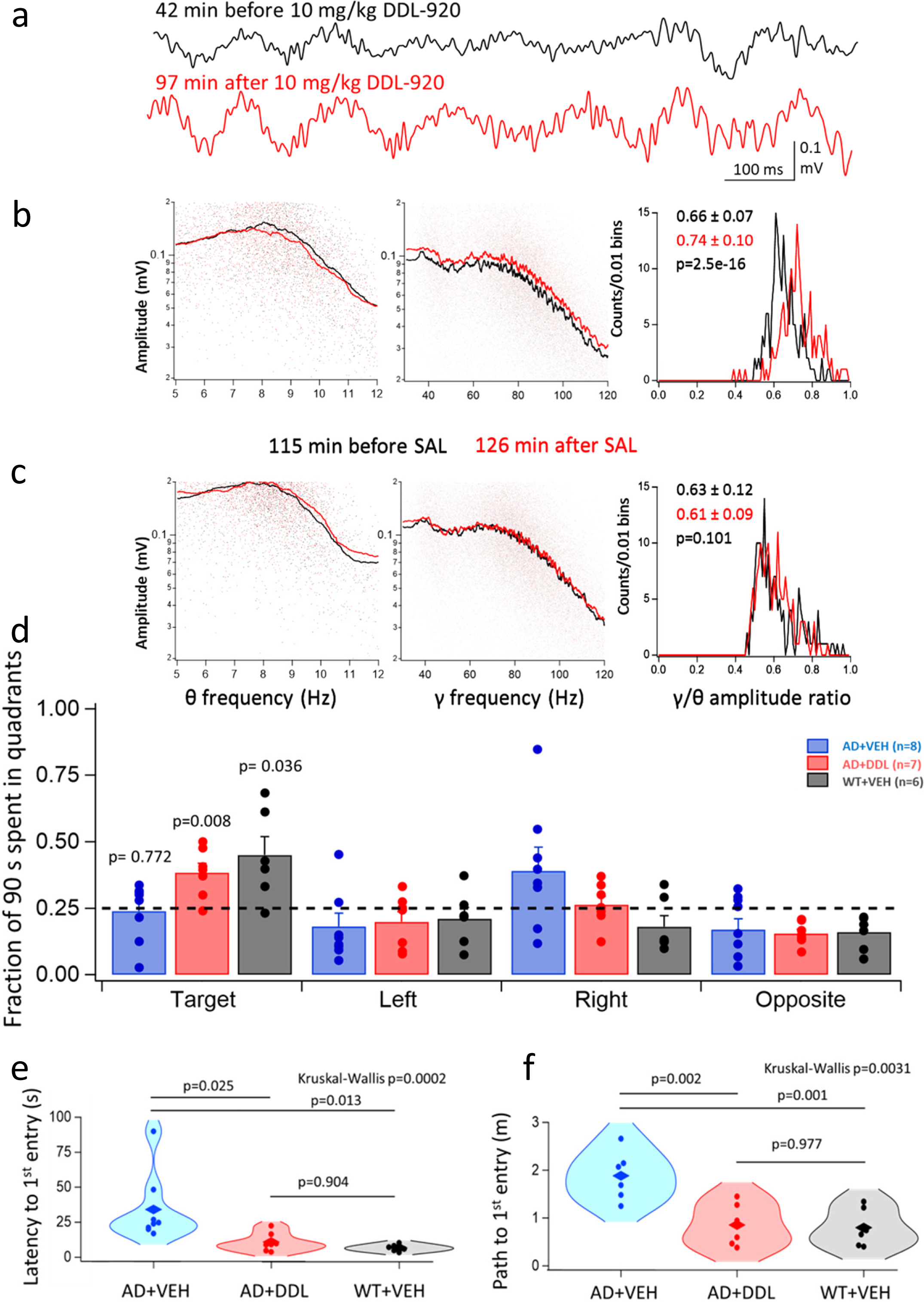
Effects of DDL-920 (10 mg/kg) oral administration with a pipettor on θ- and γ-oscillations recorded in the hippocampi of AD model mice and on their memory test in the Barnes maze. **a,** Epochs (1 s long) of *in vivo* hippocampal recordings of θ and γ-oscillations in an AD model mouse at the indicated times before (*black*) and after (*red*) the oral administration of 10 mg/kg DDL-920. **b,** Instantaneous amplitude vs frequency plots of the θ-(*left panel*) and γ-oscillations (*middle panel*) in the same mouse obtained from 180 s long epochs at the same times (*black* and *red*) as in **a**. *Right panel*: Histograms of the 180 measurements over as many s of the ratios of the averaged γ to θ oscillation amplitudes (p-values are from t-tests). **c,** Same plots as in **b**. for the same mouse after receiving a saline (SAL) oral administration one week after the injection in **a**. Color codes refer to the times before and after SAL administrations. Note the lack of differences between the averaged γ to θ oscillation amplitude ratios after SAL administrations. **d-f**: After two weeks of twice-a-day oral DDL-920 (10 mg/kg) or vehicle (VEH) administration, 15 AD mice and 6 WT animals were subjected to a one-week training and probe trials in the Barnes maze while still receiving once daily the DDL-920 or SAL oral administration (please see Supplementary Figure S9 for details). **d**, Plot of the fractions of time of the 90 s total time spent in the four quadrants of the Barnes maze (Supplementary Figure S5) on probe day 48 hrs after the last training session. The target quadrant is the quarter area of the maze that previously contained the escape hole. Vehicle (VEH) treated WT mice and DDL-920-treated AD mice spent significantly (probabilities indicated on the graph are from a t-test comparing the group means to a mean value of 0.25) more time than by chance (0.25) in the target quadrant, whereas the VEH-treated AD mice did no better than chance in this quadrant (p-values indicated on the graph are from a t-test comparing the group means to a mean value of 0.25). In addition, a non-parametric Kruskal-Wallis test (p=0.024) and a one-way ANOVA (p=0.016) indicate significant differences between the three groups in the target quadrant. Moreover, the multiple comparison Tukey test showed that the WT VEH group was significantly different from the AD VEH group (p=0.018), but not from the AD DDL-920 treated group (p=0.642), which in turn was also significantly different from the AD VEH group (p=0.042). **e**, Violin plots of the latencies of the first entries to the escape holes on the 48 hrs probe day after the training has been concluded. Individual data points are shown as solid circles and the mean values are depicted by a rhomboid symbol. The non-parametric Kruskal-Wallis test indicates that at least two groups are significantly different, and the post-hoc non-parametric Tukey test comparing all group means shows that the AD+VEH group is significantly different from both other groups, whereas the AD+DDL and WT+VEH group means are not different from each other (p-values are indicated on the graph). **f**, Violin plots of the path lengths until the first entries to the escape holes on the 48 hrs probe day. Individual data points are shown as solid circles and the mean values are depicted by a rhomboid symbol. The non-parametric Kruskal-Wallis test indicates that at least two groups are significantly different, and the post-hoc non-parametric Tukey test comparing all group means shows that the AD+VEH group is significantly different from both other groups, whereas the AD+DDL and WT+VEH group means are not different from each other (p-values are indicated on the graph).

Next, we wanted to know if the increase in γ-oscillation power produced by oral administration of DDL-920 in AD mice is beneficial to ameliorate the cognitive deficits reported in AD mouse models. Our AD model mice were 3-month-old at the beginning of the experiments ApoE4-TR:5xFAD mice (see Methods for details), as 5xFAD mice of similar age have been shown to be considerably impaired on the memory task in the Barnes maze.^71^ For two weeks, and twice a day we treated by oral administration through a pipettor 8 AD mice with vehicle, 7 AD mice with 10 mg/kg DDL-920, and 6 WT mice with vehicle. The treatment schedule and the Barnes maze testing timeline are shown in Supplementary Figure S9. By the probe testing day, the animals were 3.75-month-old. Following treatment, we measured the time spent in the quadrant of the maze containing the escape hole on the 48 hr probe day. Figure 4d illustrates that the DDL-920 treated AD mice and the saline-treated WT mice were significantly more likely than the ratio of 0.25 (chance only; statistical tests and p-values are given in the legend) to spend time in the quadrant where the escape hole was previously located. In addition, the vehicle-treated AD mice had significantly longer latencies (Figure 4e) to find the escape hole (mean ± SEM: 34.04 ± 8.68 s; median: 24.35 s; n=8 mice) than the DDL-920 treated AD mice (10.49 ± 2.41 s; median: 8.9 s; n=7 mice) or the vehicle-treated WT mice of similar age (6.72 ± 0.96 s; median: 6.65 s; n=6 mice). The non-parametric Kruskal-Wallis test indicates that at least two of the three groups are significantly different from each other (p=0.0002), and a multiple comparison Tukey test shows that the vehicle-treated AD group is significantly different from the DDL-treated AD group (p=0.025) as well as the vehicle-treated WT group (p=0.013), whereas the DDL-treated AD group is not significantly different from the vehicle-treated WT group (p=0.904; Figure 4e). In the Barnes maze, latency may be regarded as an unreliable measure because it depends on locomotion speed. Therefore, we also compared the path lengths of the animals to the target hole, as this parameter is independent of motor effects (Figure 4f). Vehicle-treated AD mice exhibited significantly longer path lengths (Figure 4f) before finding the escape hole (mean ± SEM: 1.88 ± 0.21 m; median: 1.88 m; n=8 mice) than the DDL-920 treated AD mice (0.82 ± 0.17 m; median: 0.75 m; n=7 mice) or the vehicle-treated WT mice of similar age (0.77 ± 0.16 m; median: 0.68 m; n=6 mice). For the path lengths, the non-parametric Kruskal-Wallis test indicates that at least two groups are significantly (p=0.0031) different from each other, and a multiple comparison Tukey test shows that the vehicle-treated AD group is significantly different from the DDL-treated AD group (p=0.002) as well as the vehicle-treated WT group (p=0.001), whereas the DDL-treated AD group is not significantly different from the vehicle-treated WT group (p=0.977; Figure 4f). We also plotted the total distance run by the mice during the training period and the probe trials and found no differences between the three groups (Supplementary Figure S10), indicating long-term DDL-920 treatment caused no side effects affecting motor function while neither weight loss nor adverse events were observed.

## Discussion

We describe a unique pharmacological approach meant to enhance cognitive performance and working memory in a state-dependent manner by engaging and amplifying the brain’s intrinsic γ-oscillations by enhancing the function of PV+INs. We anatomically and pharmacologically identified GABA_A_Rs assembled of α1β2δ subunits responsible for the tonic inhibition of PV+INs. We further demonstrated that DDL-920, a small molecule NAM of these receptors, is a potent, efficacious, and selective blocker of the tonic inhibition of PV+INs and consequently enhances γ-oscillations both *in vitro* and *in vivo*. In spite of the increase in γ-oscillation power, the PAC of θ- and γ-oscillations was not altered by DDL-920, which is important for working memory performance. When administered twice daily for two weeks, DDL-920 restored the cognitive/memory impairments of 3-month-old AD model mice as measured by their performance in the Barnes maze. Taken together, our findings indicate that the unique subunit composition of extrasynaptic GABA_A_Rs of PV+INs should be a valid target for boosting γ-oscillations that could be beneficial in a variety of neurological and psychiatric disorders, including AD.

A remarkable study^25^ was the first to describe that optogenetic stimulation of PV+INs or visual stimulation at a characteristic γ-oscillation frequency of 40 Hz reduced the levels of Aβ40/42 in 5XFAD mice, presumably through the involvement of microglial intermediaries. Since the publication of that study a vast amount of interest has been sparked in adapting the 40 Hz stimulation paradigm to humans^28^ for cognitive enhancement therapies, particularly in AD. The non-invasive Gamma ENtrainment Using Sensory stimulation (GENUS) protocol administered daily at 40 Hz for three months has been recently reported to partially benefit early-stage AD patients in a Phase 1 feasibility study.^72^ In spite of the successful application of various modalities of 40 Hz stimulation to humans, several studies in animals^30–32^ have shown that the exogenous 40 Hz stimulation does not entrain endogenous γ-oscillations in the brain, and in some instances it may actually interfere with the endogenous rhythms. In fact, a recent study^32^ found that 40 Hz visual stimulation was effective in reducing the Aβ plaque load in only 1 out of 6 cohorts of AD model mice. In contrast to imposing an exogenous fixed 40 Hz frequency onto the brain, the endogenous γ-rhythms of the brain may occur at various frequencies for a given task. Our approach enhances the power of innate γ-oscillations over all the frequencies (30-120 Hz) in the characteristic spectrum (e.g., Figure 3f). This should ensure that γ-oscillation power will be enhanced regardless of the frequencies demanded by the cognitive or memory tasks in the brain. The mechanism whereby DDL-920 enhances γ-oscillations is probably through increasing the gain of the rhythmic synaptic transmission onto the PV+INs required for these oscillations, both excitatory from principal cells, and inhibitory, from other PV+INs. Tonic inhibition has been identified as a powerful modulator of synaptic gain and function ^73^, and the synaptic responses of PV+INs are particularly sensitive to changes in the tonic GABA conductance. ^74^ Another possible mechanism to increase γ-oscillation power through regulation of PV+IN function is to increase their firing. Yet, a compound known to increase action potential firing rate in PV+INs by modulating the voltage-dependence of Kv3.1/Kv3.2 K^+^ channels was ineffective in increasing γ-oscillations in slices; although it did enhance these oscillations once they were dampened by prior administration of Aβ peptides. ^75^ Thus, it is tempting to conclude that selectively reducing the GABA tone in PV+INs may be one of the most effective means of increasing the power of endogenous γ-oscillations in the brain.

An important question remains the applicability of our study to humans. Do PV+INs in the human brain possess the same GABA_A_R assemblies as we described here for mice? Although there are no direct studies demonstrating this, the α1 subunits are abundant in PV+INs in both macaques^76^ and humans (Zs. Maglóczky and K. Tóth, Institute of Experimental Medicine, Budapest, Hungary, personal communication). The α1 subunits’ close partnership with δ subunits in human PV+INs also awaits direct demonstration, but this dual subunit assembly is highly evident in schizophrenic patients, where α1 and δ GABA_A_R subunit mRNA changes are tightly coupled when PV+IN deficits are prevalent.^77^ There are no specific drugs known to enhance γ-oscillations in humans, but a learned compassion meditation state has been reported to significantly enhance these oscillations.^78^ It remains to be determined whether trained and frequent practitioners of this type of meditation are less affected by AD.

We are encouraged by the lack of obvious side effects of DDL-920, such as potentially inducing abnormal excitability in the brain, as after all, it is a NAM of inhibitory GABA_A_Rs. Moreover, we observed no hyperactivity, abnormal behavior, or visible side effects following its long-term (3 weeks) administration. More studies will be needed to establish DDL-920’s target selectivity, potential toxicity, and off-target effects, but our results so far are promising, and show DDL-920 has good drug-like properties, including stability in human plasma, microsomal stability, and kinetic water solubility. Our continuing drug development based on DDL-920 as a lead compound will involve additional preclinical testing in other murine models, analysis of AD related biomarkers and any off-target effects in the brain along with IND-enabling studies. We are aware of the drawbacks of the applicability of animal models of complex neurodegenerative disorders such as AD to the human condition.^79–81^ However, at the same time, we are optimistic that a potent and selective orally active small molecule enhancer of γ-oscillations should be beneficial not only in AD especially during its early stages, but in several neurological and psychiatric disorders including recovery from stroke or post-traumatic brain injury, schizophrenia, depression, ASD, and perhaps also in normal aging-related cognitive deficits.

## Supporting information

Manuscript with figures

## Acknowledgements

Parts of this research were supported by NIH/NIA grant R01AG050474 and the Coelho Endowment to I.M., NIH/NINDS Grant R01NS102608 to C.R.H. We are indebted to Dr. Keith Vossel, Director of the Mary S. Easton Center for Alzheimer’s Disease Research and Care who provided funds for research from various anonymous donors to the Center. We thank Professor Greg Cole, UCLA Department of Neurology, for providing the initial breeders of the ApoE4-TR:5xFAD mice that was used to generate the cohort for the in vivo studies with DDL-920. We would like to thank W. Sieghart (Medical University of Vienna, Austria) and J.-M Fritschy (Univ. Zürich, Switzerland) for generously sharing their GABA_A_R subunit specific antisera.

## Author Contributions

I.M. conceived the utility of a NAM specific for the GABA_A_Rs of PV+INs to enhance γ-oscillations, analyzed data, prepared figures, and wrote the initial draft of the manuscript. V.J. lead the medicinal chemistry and supervised members of the DDL. C.R.H. lead and supervised anatomical studies. X.W. and J.J.C. designed and performed experiments, analyzed data, prepared figures, and wrote parts of the methods. B.J. performed molecular simulations. D.W., W.C., J.L., C.Z., and C.S.H. carried out various experiments. L.M. analyzed data and provided software analytical procedures. All authors commented on the manuscript.

## Methods

### Key resources table

| REAGENT or RESOURCE | SOURCE | IDENTIFIER |
| --- | --- | --- |
| **Antibodies** |  |  |
| Anti-GABA(A)R alpha 1 subunit (guinea pig) | H. Mohler and J.-M. Fritschy, University of Zurich; Zurich; Switzerland | — |
| Anti-GABA(A)R alpha 1 subunit (extracellular) (guinea pig) | Alomone Labs Cat# AGP-083-GP | RRID:AB_2756618 |
| Anti-GABA(A)R beta 2 subunit (rabbit) | PhosphoSolutions Cat# 862-GB2C | RRID:AB_2492109 |
| Anti-GABA(A)R beta3 subunit (rabbit) | Millipore Cat# AB5563 | RRID:AB_177525 |
| Anti-GABA(A)R delta subunit (rabbit) | W. Sieghart, Center for Brain Research, Medical University of Vienna; Vienna; Austria. (Delta (17-60)) | RRID:AB_2827804 |
| Anti-Parvalbumin (rabbit) | K.G. Baimbridge, University of British Columbia; British Columbia; Canada (R301) | RRID:AB_2315236 |
| Anti-Parvalbumin (mouse) | Millipore Cat# MAB1572 | RRID:AB_2174013 |
| Goat anti-Rabbit IgG (H+L) Highly Cross-Adsorbed Secondary Antibody, Alexa Fluor™ Plus 488 | Thermo Fisher Scientific Cat# A32731 | RRID:AB_2633280 |
| Goat anti-Guinea Pig IgG (H+L) Highly Cross-Adsorbed Secondary Antibody, Alexa Fluor™ 488 | Thermo Fisher Scientific Cat# A-11073 | RRID:AB_2534117 |
| Goat anti-Mouse IgG (H+L) Highly Cross-Adsorbed Secondary Antibody, Alexa Fluor™ 488 | Thermo Fisher Scientific Cat# A-11029 | RRID:AB_2534088 |
| Goat anti-Guinea Pig IgG (H+L) Highly Cross-Adsorbed Secondary Antibody, Alexa Fluor™ 555 | Thermo Fisher Scientific Cat# A-21435 | RRID:AB_2535856 |
| Goat anti-Rabbit IgG (H+L) Highly Cross-Adsorbed Secondary Antibody, Alexa Fluor™ Plus 555 | Thermo Fisher Scientific Cat# A32732 | RRID:AB_2633281 |
| **Biological samples** |  |  |
| Human liver microsomes | Thermo Fisher Scientific | HMMCPL |
| Blood plasma | Stemcell Technologies Inc | 70039 |
| **Chemicals, peptides, and recombinant proteins** |  |  |
| NADPH | Millipore Sigma | 10107824001 |
| MgCl<sub>2</sub> | Millipore Sigma | M4880-100G |
| Acetonitrile | Thermo Fisher Scientific | A998-4 |
| DMSO | Thermo Fisher Scientific | BP231-1 |
| ProLong. Fluoromount G Mounting Medium | Thermo Fisher Scientific. Catalog number. P36934 | RRID:SCR_015961 |
| DDL-920 | UCLA DDL | Ref XX |
| **Critical commercial assays** |  |  |
| CHIRALPAK® CAN | Chiral Technologies | H15L-109 |
| IAM.PC.DD | Regis Technologies | 774011 |
| Micro BCA™ Protein Assay Kit | Thermo Fisher Scientific | 23235 |
| Slide-A-Lyzer™ MINI Dialysis Devices | Thermo Fisher Scientific | PI88401 |
| **Deposited data** |  |  |
| β3 homopentameric GABA(A)R | RCSB Protein Data Bank (RCSB PDB) | 4COF |
| β3δ heteropentameric GABA(A)R | RCSB Protein Data Bank (RCSB PDB) | 7QND |
| **Experimental models:Mus musculus** |  |  |
| C57BL/6J | Jackson Laboratory | RRID:IMSR_JAX:000664 |
| Gabrd-/- mice on a C57BL/6J background | Jackson Laboratory | RRID:IMSR_JAX:003725 |
| Pvalb-IRES-Cre mice | Jackson Laboratory | RRID:IMSR_JAX:017320 |
| B6.Cg-Gt(ROSA)26Sortm14(CAG-tdTomato)Hze/J | Jackson Laboratory | RRID:IMSR_JAX:007914 |
| ApoE4-TR:5xFAD mice | Greg Cole (UCLA) |  |
| **Software and algorithms** |  |  |
| ANY-maze software | Stoelting Co | RRID:SCR_014289 |
| Cresset software | Flare v7.2 |  |
| Optibrium software | StarDrop 7.5 |  |
| Igor Pro 6.32, 7, 8, 9 | Wavemetrics | RRID:SCR_000325 |
| iSpy video motion analysis software | https://www.ispyconnect.com/ |  |
| ZEN Digital Imaging for Light Microscopy | Carl Zeiss light microscopy imaging systems | RRID:SCR_013672 |

## Resource availability

### Lead contact

Further information and requests for resources and reagents should be directed to and will be fulfilled by the lead contact, Istvan Mody.

### Materials availability

All new material and mouse models generated from this study will be available upon request. Some may require MTAs.

### Data and code availability

- All data will be made available upon request from the lead author through Dryad.
- Custom Igor procedures are available from the lead contact upon request.
- Any additional information required to reanalyze the data reported in this work paper is available from the lead contact upon request.

### Experimental model and study participant details

All procedures were performed in accordance with protocols approved by the UCLA Institutional Animal Care and Use Committee (IACUC) and guidelines of the National Institutes of Health. Mouse strains used were C57BL/6J (Black 6, Jackson Laboratory; JAX stock #000564), JAX stock #003725 (Gabrd-/- mice on a C57BL/6J background), PVcre/Ai14 (JAX stock#017320/#007914). All non-AD mice were 4-5 months-old at the time of the experiments. The AD model mice consisted of ApoE-TR mice which express APOE4 under the control of the endogenous mouse APOE promoter were bred to 5xFAD mice (Tg6799) which co-express five FAD mutations (APP K670N/M671L + I716V + V717I and PS1 M146L + L286V) under the control of the neuron-specific mouse Thy-1 promoter, and backcrossed three times to ApoE-TR mice, resulting in mice homozygous for APOE4, and hemizygous for the 5XFAD transgenes, on a background strain 97% C57Bl/6J and 3% SJL; mice were then inbred between 5xFAD+ and 5xFAD-resulting in littermates E4+/+:FAD+ and E4+/+:FAD-. The AD model mice were 4-5 months old for the in vitro and in vivo electrophysiological recordings, and 3 months-old when training for the Barnes maze experiments were started.

### Method details

#### Tissue collection for light microscopy

For immunohistochemistry studies, adult mice were deeply anesthetized with Fatal-Plus (90 mg/kg, i.p.) and perfused transcardially with 4% paraformaldehyde in 0.1M phosphate buffer (pH 7.3). Brains were removed and postfixed for 1 h in the same fixative. Following cryoprotection, forebrain regions containing the hippocampus were frozen and sectioned coronally or sagittally at 30 µm with a cryostat. Free-floating sections were stored in cryoprotectant for subsequent immunohistochemical studies.

#### Immunohistochemistry for light microscopy

Tissue processing and immunohistochemistry were performed with previously published methods^82^. For double immunofluorescence labeling, free-floating sections (30 µm) were incubated for 72 h at room temperature in the following combinations of primary antisera: 1) rabbit anti-parvalbumin (1:5,000; KG Baimbridge, R301) and either guinea pig anti-α1 subunit (1:30,000; J-M Fritschy) or guinea pig anti-α1 subunit (1:2,000; Alomone, AGA-001-GP); 2) rabbit anti-δ subunit (1:3,000; W. Sieghart) and either mouse anti-parvalbumin (1:2,000; Millipore, MAB1572) or guinea pig anti-α1 subunit (J-M Fritschy, as above); 3) guinea pig anti-α1 subunit (Alomone, as above) and either rabbit anti-β2 subunit (1:100; PhosphoSolutions, 862-GB2C) or anti-β3 subunit (1:250; Millipore, AB5563). After thorough rinsing, sections were incubated with a combination of fluorescence-labeled secondary antisera (1:500), either 1) goat anti-rabbit IgG conjugated to Alexa Fluor Plus 488 (Thermo Fisher Scientific, A32731) and goat anti-guinea pig IgG conjugated to Alexa Fluor 555 (Thermo Fisher Scientific, A21435); or 2) goat anti-rabbit IgG conjugated to Alexa Fluor Plus 555 (Thermo Fisher Scientific, A32732) and either goat anti-guinea pig IgG conjugated to Alexa Fluor 488 (Thermo Fisher Scientific, A11073) or goat anti-mouse IgG conjugated to Alexa Fluor 488 (Thermo Fisher Scientific, A11029). Sections were incubated in secondary antisera for 4 h at room temperature, mounted on slides and cover slipped with anti-fade medium ProLong Diamond (Thermo Fisher Scientific).

Fluorescence-labeled sections were scanned with an LSM 880 confocal microscope (Carl Zeiss), and Z-stack images (0.2 or 1.0 µm optical thickness) were acquired (excitation spectra 488 and 555nm) with 20x and 63x objectives. For double-labeled sections, optical slices were scanned separately for each label, alternating between the two channels in each section through the Z-stack. Confocal images were analyzed with Zen 2 software (Carl Zeiss).

#### Electrodes used for in vivo recordings

Electrodes (Plastics One) were stainless-steel, with polyimide electrode insulation, ending in a socket that fitted into the custom-made recording system. Electrode lengths (measured from bottom of socket contact to tip of wire) were 5 mm (for the two unilaterally implanted electrodes) and 25 mm for the ground/reference electrode. Electrode diameters were 125 μm bare and 150 μm insulated. The two unilaterally implanted electrodes were custom made as follows: a socket (gold-plated stainless steel Amphenol contact, O.D: 1.07 mm, I.D: 0.686 mm and length: 7.87 mm) was attached to a flexible 10 mm long PFA Teflon insulated wire. The free tip of this wire was soldered to another 3 mm long Amphenol socket attached to the 5 mm electrode. The openings of the sockets were trimmed to 1 mm length, cleaned from debris, smoothened on the edges and tightened. The three electrodes were assembled into one unit by connecting their sockets with J-B Weld^TM^ two-part epoxy. The space between the sockets corresponded to the distance between the three Amphenol pins soldered to the recording preamplifier.

#### Stereotaxic surgeries

The two recording electrodes were implanted bilaterally in the ventral hippocampal CA1 region at −3.3 AP, 3.5 ML, 3 DV (AP, anterior-posterior; ML, medial-lateral; DV, dorsal-ventral) according to established coordinates^83^ and from the ALLEN Mouse Brain Atlas, Version 2 (2011), Allen Institute for Brain Science. These coordinates consistently positioned the electrodes in the pyramidal cell layer or slightly above it, as judged by the depth profile of the relationship between θ phase and γ oscillations^84^ in the shape of the recorded SPW-R complexes^66,85^. The reference/ground electrode was implanted over the cerebellum. The skull and the lower part of the electrode unit system was then covered with Ortho-Jet^TM^ acrylic resin. During the surgery, lidocaine (2 µl from 2% solution) was injected subcutaneously on the neck and the nonsteroidal anti-inflammatory drug Rimadyl (Carprofen) (0.1 mg/kg) was administered intraperitoneally for pain management (Rimadyl was additionally administered for two consecutive days). In addition, 50 µl sterile saline was administered subcutaneously for hydration during surgery. During recovery after surgery (one week) mice were single housed in plastic disposable cages with a plastic lid that did not interfere with the head implants.

#### In vivo recordings and data acquisition

The electrical signals from the mouse brain were relayed through a custom-made recording system to a computer, located outside the recording room. Briefly 1-1.5 m long cable was constructed using 8 intertwined bare copper wires (single 8058-Magnet Wire) and placed inside a PVC tubing. The cable was soldered accordingly to a LT1112S8#PBF General Purpose Amplifier 2 Circuit 8-SO (Digi-Key) on one end and to a MMA25-011 connector plug with male pins (Digi-Key) to the other end. The connector plug end of the cable was attached to a 10-channel slip ring commutator (Campden Instruments). Both ends of the cable were covered with J-B Weld^TM^ two-part epoxy. The commutator was joined to a custom-made connector box that also contained two 9 V batteries providing the power supply to the preamplifier. The signal was then transmitted to Model 3600 16 channel amplifier (A-M Systems) (LP: 300 Hz, HP: 0.3 Hz Gain:1K). The output signal was led to an USB 6009 14 bit A/D converter (National Instruments) connected to a laptop computer. Igor 8.0 (Wavemetrics) NIDAQ tools was used to record the two channels in each mouse at a sampling rate of 2048 s^-1^. During the continuous recording sessions the mice were housed in clear polycarbonate containers (32 cm diameter and 38 cm high, Cambro) with the floor covered by bedding. Food, water, and nest material were also added. The video rate captured by an infrared-sensitive USB camera was 11-13 frames/s providing a movement resolution of ∼76-90 ms. The iSpy software was used to determine the motion of the animals through a frame-by-frame subtraction of successive video images. The threshold for movement detection of the software was set to 80 pixels.

#### Analysis of in vivo LFP recordings

All data analyses were carried out using custom written procedures in IgorPro 8.0 and 9.0 (Wavemetrics) using its built-in functions for RMS measurement, FIR filtering, FFT, Hilbert amplitude, etc. The rest of the analyses such as Morlet wavelet transform, spectral peak detection, PAC, and FAC were done using procedures written in Igor 8.0 based on published methods. First, using a FIR filter (using at least 401 coefficients or with the number of coefficients determined as = int(50/(22*(HF-LF)/SR)), where int is the integer part, 50 is the cutoff of the filter in dB, HF and LF are the high and low frequencies, respectively of the bandpass, and SR is the sampling rate (2048 s^-1^). We bandpass filtered the raw recordings, at 0.5-4 Hz for δ, 5-12 Hz for θ and 30-120 Hz for γ oscillations. This digital FIR filtering did not change the phase of the oscillations^66^. The band-pass filtered traces were then used for calculating the RMS values for the analyses.

Several recording segments of 180 s duration were selected during intervals of 3 hrs before and within 3 hrs after the administration of drug or vehicle. The criterion for selection was based on the RMS of θ-oscillations that had to be larger than the mean + 3xSD of the average RMS of the θ-filtered recording for the entire 12 hr period. The choice of the θ-oscillations for the selection was to avoid bias by selecting segments based on the power of γ-oscillations. One segment pre- and one after the drug or vehicle administration was each selected randomly from those that satisfied the selection criterion. We used published methods^68,69,86,87^ to calculate the modulation indices of γ oscillations by θ oscillation frequency or phase. We used two methods to determine the coupling of θ oscillation phase to γ oscillation amplitude. In the first method we started by isolating the individual θ cycles from the θ-filtered traces during the 8 s epochs yielding about 50-70 cycles in total. Then, each θ cycle was divided into equal time segments each corresponding to π/10 radians of the given θ cycle. The γ bandwidth filtered traces were then aligned with these time segments and their peak-to-peak amplitudes and frequencies were measured. The median values of these amplitudes were entered into a matrix in the corresponding γ-frequency row binned by 10 Hz from 30-120 Hz. The matrix was complete when all the θ cycles during the 8 s epoch were analyzed. The second method used the Hilbert transform of the θ bandwidth filtered signal over the 8 s epoch to calculate the continuous phase of the θ oscillations. Over each phase of the θ cycle thus obtained, the raw signals were filtered in 10 Hz bins between 30-120 Hz (9 bins in total) and the amplitude histograms of these band-filtered signals were plotted over each θ phase. The two methods yielded nearly identical PAC results, but we decided to show the graphs obtained by the first method. Once the various modulation matrices were obtained for at least 10 different 8 s periods during the same chosen 180 s epoch, we averaged the matrices. The resulting averages were converted to a Z-scale (Figure 3d) for every 10 Hz of the γ-frequency band and were smoothed by a 2D spline algorithm (ImageInterpolate function of Igor 8.0). The spectrograms of the long-duration recordings were obtained by using the Short-Time Fourier Transform (STFT) function of Igor Pro 8.0 or 9.0 with using a Hanning window with a segments size of 60 s (122,880 points) and a 30 s overlap.

#### Slice preparation

Mice were at least 3-months-old when the ex vivo experiments were undertaken. They were anesthetized with isoflurane and decapitated following UCLA Chancellor’s Animal Research Committee protocol. Horizontal 350 μm thick slices were cut on a Leica VT1200S vibratome in ice-cold N-Methyl-D-Glutamine (NMDG)-based HEPES-buffered solution, containing (in mM): 135 NMDG, 10 D-glucose, 4 MgCl_2_, 0.5 CaCl_2_, 1 KCl, 1.2 KH_2_PO_4_, 20 HEPES, 27 sucrose (bubbled with 100% O_2_, pH 7.4, 290-300 mOsm/L). Then, slices were incubated at 32°C in a reduced sodium artificial CSF (ACSF), containing (in mM): NaCl 85, D-glucose 25, sucrose 55, KCl 2.5, NaH_2_PO_4_ 1.25, CaCl_2_ 0.5, MgCl_2_ 4, NaHCO_3_ 26, pH 7.3-7.4 when bubbled with 95% O_2_, 5% CO_2_. After 30 min low sodium ACSF was substituted for normal ACSF at room temperature, containing (in mM): NaCl 126, D-glucose 10, MgCl_2_ 2, CaCl_2_ 2, KCl 2.5, NaH_2_PO_4_ 1.25, Na Pyruvate 1.5, L-Glutamine 1, NaHCO_3_ 26, pH 7.3-7.4 when bubbled with 95% O_2_, 5% CO_2_. All salts were purchased from Sigma-Aldrich.

#### In vitro gamma oscillations

Kainic acid (KA, Tocris) was used to generate gamma γ-oscillations in slices. To get a stable level of γ-oscillations after the slice cutting and recovery incubation period, the slices were incubated for at least 30 min in ACSF containing 50 nM KA before transferring them to the recording chamber^40,42^. Recordings were done in an interface chamber at 34°C perfused with normal ACSF also containing 50 nM KA at a speed of 5 ml/min. Oscillatory network activity was recorded in CA3 stratum pyramidale with the use of a patch pipette (3-5 MΩ resistance) filled with KA-containing ACSF connected to the headstage of an amplifier (A-M Systems Inc., model 3000). The signal was band-pass filtered between 0.1 and 1000 Hz and then was fed through an instrumentation amplifier (Brownlee BP Precision, model 210A) and sampled at 4096 s^-1^ with a National Instruments A/D board. Field potentials were recorded using a custom LabView software (EVAN) and analyzed with a custom written procedure (Wavemetrics, IGOR Pro 8). Peak frequencies, power at peak frequency and total power were obtained from the corresponding RMS, averaged from 60 s recording periods.

#### Patch clamp recordings and analyses

For recording, brain slices were transferred to a submerged recording chamber at 34°C and perfused at 5 ml/min with ACSF. Slices were visualized under IR-DIC upright microscope (Olympus BX-51WI, 20x XLUMPlan FL N objective) and whole-cell recordings were obtained from either dentate gyrus granule cells, somatosensory pyramidal cells, or PV+INs labeled with tdTomato with borosilicate patch pipettes (4-6 MΩ, King Precision Glass) containing of internal solutions (ICS) (in mM): 140 Cs-met, 2 MgCl_2_ 10 HEPES, 0.2 EGTA, 2 Na_2_-ATP, 0.2 Na_2_-GTP.

The pH of the ICS was adjusted to 7.2 with CsOH and its osmolarity was 285-290 mOsm. ICS were stored at −80°C in 1 ml aliquots. Before each experiment, ICS aliquots were thawed to room temperature and kept on ice during recording. Recordings were obtained using an Axon-patch 200B amplifier (Molecular Devices, San Jose, CA, USA), low-pass filtered at 5 kHz (Bessel, 8-pole) and digitized at 10 kHz with a National Instruments data acquisition board (BNC 2110, National Instruments, Austin, TX, USA). All data were acquired with EVAN (custom-designed LabView-based software). Spontaneous inhibitory post synaptic currents (sIPSCs) were recorded at a holding potential (Vh) of 0 mV. Whole-cell capacitance were estimated from fast transients evoked by a 5 mV voltage command step using lag values of 7 μs and then compensated to 70-80%. The series resistance was monitored before and after the recording, recordings with series resistances >20 MΩ or a change >20% during the recording were excluded.

A custom written procedure in IGOR Pro 8.0 (Wavemetrics) was used to perform the analysis of tonic and phasic currents^88^. An all-points histogram of a randomly selected recording segment of 30 s during the period of interest was plotted. A Gaussian was fitted to the part of the distribution from the minimum value at the left to the rightmost (largest) value of the histogram distribution. The mean value of the fitted Gaussian was taken as the tonic current (I_tonic_). The skewed distribution toward synaptic events (I_phasic_). This process was repeated for all segments of interest.

#### In Silico Screening

There is no crystal or cryo-EM structural study of the α1β2δ GABA_A_R complex. This is available for the β3 homopentameric and the β3δ heteropentameric GABA_A_R, the β2 and β3 receptors have a sequence similarity of >95% in their ligand binding and membrane spanning domains^89^. Therefore, we used it in our in silico screening. Small molecules of various structural classes were docked into the model prepared from the known crystal structure of the β3 homopentameric GABAAR (pdb:4COF)^13^. Docking of the structures was performed using Flare v7.2 (Cresset software) and the binding score of the docked structure in the model was determined. This analysis led to the identification of DDL-920 that showed good binding with a Lead Finder (LF) rank score of −10.35 in the docking simulation (Extended Data Fig. 6a). To determine if DDL-920 would bind and interact with a GABA_A_R containing the δ-subunit we used the published β3δ heteropentameric GABA_A_R structure (pdb:7QND)^14^. The docking simulation with DDL-920 and structure pdb:7QND reveals that in the presence of the δ-subunit it reaches a better LF rank score of −11.21 compared to docking to pdb:4COF. The model also reveals that DDL-920 makes key interactions with both the β3 and δ subunits of pdb:7QND (Extended Data Fig. 6b). Additionally, predictive analysis for brain permeability and other drug-like properties was done *in silico* using StarDrop (Optibrium software) analysis platform. The analysis showed that DDL-920 would be brain permeable and with good drug-like properties. Hence it was selected for synthesis and testing in our *in vitro* assays and *in vivo* model.

#### Pharmacokinetics (PK) in mice

All *in vivo* experiments described were carried out in strict accordance with good animal practice according to NIH recommendations. All procedures for animal use were approved by the Animal Research Committee (ARC) at UCLA and under an approved ARC protocol. Wildtype (WT) C57Bl6 mice were dosed by oral (gavage or pipette) or subcutaneous (SC) injection at 3 doses (1, 5, 10 mg/kg), and animals were euthanized 1, 3 and 6 hours after administration by ketamine/xylazine over-anesthesia following by transcardial collection of blood for isolation of plasma and saline perfusion. Brain tissue was collected *post-mortem* for assessment of compound levels.

Analysis of brain concentrations was done at the UCLA Pasarow Mass Spectrometry Lab (PMSL; Kym Faull, Ph.D., Director). A targeted liquid chromatography-tandem mass spectrometry (LC-MS/MS) assay was developed for DDL-920 using the multiple reaction monitoring (MRM) acquisition method on a 6460 triple quadrupole mass spectrometer (Agilent Technologies) coupled to a 1290 Infinity HPLC system (Agilent Technologies) with a Phenomenex analytical column (Gemini-NX 3µm C18 110 Å 100 x 2.0 mm). The HPLC method utilized a mixture of solvent A (99.9/1 Water/Formic Acid) and solvent B (99.9/1 Acetonitrile/Formic Acid) and a gradient was use for the elution of the compounds (min/%B: 0/0, 5/0, 20/99, 22/99, 25/0, 35/0).

In this assay, detection of a fragment ions originating from DDL-920 (m/z: 308.1 −> 141, 235) as well as LC retention time (RT= 11 min) were utilized to ensure compound specificity and accurate quantification in the biological samples. An internal standard (IS) that closely resembled DDL-920 (m/z: 361; 100 pmol; RT = 14.9) was added to every sample to account for compound loss during sample processing. Standards were made in drug naïve plasma and brain lysates with increasing amounts of cambinol (S1, S2: 0 pmol/ S3,S4: 1 pmol/ S5,S6: 10 pmol/ S7,S8: 100 pmol, S9,S10: 1000 pmol). The standard curve was made by plotting the amount of DDL-920 per standard vs. the ratio of measured chromatographic peak areas for each compound (DDL-920/IS). The trendline equation was then used to calculate the absolute concentrations of DDL-920 in the brain tissue.

#### Efficacy testing in E4FAD mice

Male and female 3-month-old mice were administered DDL-920 via pipette feeding by the method of Atcha et al^15^ at a dose of 10 mg/kg BID (20 mg/kg/day). Mice were weighed before the first dose for the calculation of volume to be administered. The formulation used comprised DDL-920 as a 10 mg/mL solution dissolve first in water and then in strawberry syrup (50 V/V final water concentration). Mice were dosed for 2 weeks BID (Supplementary Figure S9) before entering Barnes Maze testing as describe by Attar et al^16^. Briefly, in a room with visual cues on the walls, mice are introduced in a circular maze containing 20 holes where 19 holes are closed during training and all 20 holes are closed during the probe trials. On day 1 of the habituation stage, mice are placed at the center of the maze inside the start chamber. After 10 s the start chamber is removed, and mice are allowed to explore the maze for 3 min. After 3 min of exploration, the mice are guided to the target (escape) hole and allowed to stay there for 2 min. After habituation, mice undergo 3 consecutives training days, and each training day is comprised of 3 trials. Each trial starts with the mice being placed at the center of the maze inside the start chamber for 10 s. After this the start chamber is removed and mice are allowed to explore the maze for 3 min. If during this time they do not find the target hole, they are guided to it and allowed to remain inside for 1 min. At the end of each training day mice were dosed at 10 mg/kg. Next are the 24- and 48-hours probe days. During the probe days, the target hole is closed, and spatial memory of the mouse is determined during a 90 s probe trial. Mice are placed at the center of the maze inside the start chamber, after 10 s the start chamber is removed, and mice are allowed to explore the maze for 90 s. At the end of the trial, the mice are returned to the home cage and dose at 10 mg/kg. The ANY-maze software (Stoelting) was used to analyze the behavior of the mice in the Barnes Maze. Over the course of the training and probe trials, the software-calculated parameters included, but were not limited to, total distance traveled, speed, number of holes visited, latency to visiting the target/escape hole, percent time spent in the four quadrants of the maze.

#### Absorption, Distribution, Metabolism. Excretion (ADME)

Methods for determination of kinetic solubility, plasma stability, liver microsome stability, parallel artificial membrane permeability assay (PAMPA), plasma protein (human serum albumin, HSA) binding, and brain tissue binding are described next.

#### Kinetic solubility assay

DDL-920 (10 mM, 100% DMSO) was diluted separately into aqueous buffer (100 µM; PBS pH 7.4) and DMSO at various concentrations (1000, 100, 10, 1, 0.1 µM). The solutions were incubated at 37°C for 90 min and centrifuged at 16K x g for 5 min. An aliquot of each supernatant was analyzed by LC-MS/MS. A standard curve was generated by plotting the known amount of analyte per standard in DMSO vs. chromatographic peak area. Kinetic solubility (mM) was calculated using the trendline equation with maximum chromatographic peak area observed in the aqueous DDL-920 sample^17^.

#### Liver microsome stability assay

A 1 µL sample of 1 mM DDL-920 in 100% DMSO was added to 1 mL of 0.5 mg/mL human liver microsomes (Thermo Fisher Scientific, Cat# HMMCPL) in PBS pH 7.4, 2 mM NADPH, and 2 mM MgCl_2_; and incubated at 37 °C for 120 min. Aliquots (50 µL) of the microsome solution were taken at timepoints of 0, 5, 10, 15, 30, 60, 90, and 120 min, and added to a 200 µL 100% acetonitrile reaction quenching solution containing an internal standard. Solutions were clarified by centrifugation (16K x g, 5 min) and supernatants transferred to new tubes and lyophilized.

Samples were reconstituted in 100 µL of 50/50/0.1 water/acetonitrile/formic acid prior to analysis via LC-MS/MS^18^. Normalized chromatographic peak areas were plotted at each time point and the half-life (T_1/2_) of compound in liver microsomes was determined by using the trendline equation to calculate the time at which compound abundance was 50% of that at timepoint 0 (T_0_).

#### Plasma stability assay

A 1 µL DDL-920 (1 mM in 100% DMSO) was added to human peripheral blood plasma (Cat#70039, Stemcell Technologies Inc) diluted 2-fold in PBS (1000 µL, pH 7.4) and incubated at 37°C for 180 min. Aliquots (50 µL) of the plasma solution were taken at 0, 15, 30, 60, 90, 120, 150, and 180 min, and added to a 200 µL 100% acetonitrile reaction quenching solution containing an internal standard. Solutions were clarified by centrifugation (16K x g, 5 min) and supernatants were transferred to new tubes and lyophilized. Samples were reconstituted in 100 µL of 50/50/0.1 water/acetonitrile/formic acid prior to analysis via LC-MS/MS). Normalized chromatographic peak areas were plotted at each time point and the T_1/2_ in plasma was determined by using the trendline equation to calculate the time at which compound abundance was 50% of that at timepoint 0 (T_0_).

#### Human serum albumin (HSA) binding assay

The liquid chromatography-ultraviolet/visible spectroscopy (LC-UV/Vis) assay was performed on a 1290 Infinity HPLC system (Agilent Technologies) with an HPLC column containing immobilized HSA (CHIRALPAK® CAN, Chiral Technologies, Cat#H15L-109, 5 µm 50 x 4 mm). The HPLC method utilized an isocratic gradient containing 100% 6.7 mM phosphate buffer saline (pH 7.4). The retention time of the compound (tr) and injection peak were recorded.

The percent protein binding (P) was calculated using the following equations:

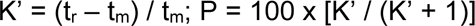

#### Parallel artificial membrane permeability assay (PAMPA)

A liquid chromatography-ultraviolet/visible spectroscopy (LC-UV/Vis) assay was performed on a 1290 Infinity HPLC system (Agilent Technologies) with an HPLC column containing immobilized phosphatidylcholine (IAM.PC.DD, Regis Technologies, Cat#774011, 5 µm 300 Å 100 x 4.6 mm). The HPLC buffer comprised 6.7 mM PBS (pH 7.4; solvent A) and acetonitrile (solvent B): a gradient was used for the elution of the compounds (min/%B: 0/20, 20/60, 21/20, 30/20). The retention time of the compound (tr) and void volume time of the column (t_0_) were recorded.

Membrane permeability (Pm) was calculated using the following equations as described^19^.

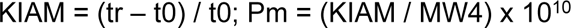

Compounds with a Pm >0.85 are predicted to be blood-brain barrier (BBB) permeable (CNS+) at pH 7.4^20^.

#### Brain Tissue Binding Assay

Brain tissue was homogenized in PBS (pH 7.4) (1: 3 weight(mg)/volume(µL)) and the protein concentration determined using the Micro BCA™ Protein Assay Kit (Thermo Fisher Scientific, Cat#23235). Brain homogenate was diluted to 20 mg/mL in PBS (pH 7.4) and added to Slide-A-Lyzer™ MINI Dialysis Devices, 10K MWCO dialysis cups (Thermo Fisher Scientific, Cat#PI88401) in a 48-well plate containing PBS (500 µL; pH 7.4). 1 µL of 1 mM compound was added to the brain homogenate (final concentration: 2 µM compound, 0.5% DMSO) and incubated on a rocker for 4.5 hours at 37 °C. 50 µL of brain homogenate (within the dialysis cup) and PBS (within the 48-well plate) were transferred to new microcentrifuge tubes containing 200 µL of quenching reagent (100% acetonitrile) containing internal standard.

Solutions were clarified by centrifugation (16,000 x g, 5 min) and the supernatants were transferred to new tubes and lyophilized. Samples were reconstituted in 100 µL of 50/50/0.1 (Water/Acetonitrile/Formic Acid) prior to analysis via LC-MS/MS. The % of the unbound drug was calculated using the following equation: f_unbound_ = [normalized peak area in buffer side / normalized peak area in brain side] x 100.

#### Quantification and statistical analysis

All statistical analyses are clearly indicated in the text or figure legends. We used the various built-in statistical functions of Igor Pro 8.0 or 9.0 for the statistical comparisons. The measure of dispersion was the SEM, unless otherwise noted. Mean values, and occasionally median values are given. The p-values for the various statistical comparisons are clearly indicated in the main text, figure legends, or in the figures themselves. The numbers of replicates are always given, and it is clearly specified what these numbers refer to.

