## Supplementary material for "A therapeutic small molecule lead enhances γ-oscillations and improves cognition/memory in Alzheimer’s disease model mice": Manuscript with figures

Department of
[1]Neurology
[2]Neurosurgery
[3]Neurology, Drug Development Laboratory, Mary S. Easton Center for Alzheimer's Disease Research
[4]Neurobiology
[6]Physiology
The David Geffen School of Medicine at UCLA, Los Angeles, CA 90095, USA

[5]Department of Electrical Engineering, Sapientia Hungarian University of Transylvania, Târgu Mureş, 540485 Romania

[‡] These authors contributed equally to this work

[†] Senior authors

* Corresponding and lead author:

Istvan Mody, Ph.D.
Department of Neurology, NRB1 Rm 575D
The David Geffen School of Medicine at UCLA
635 Charles Young Drive South
Los Angeles, CA 90095

ORCID ID: 0000-0002-7975-7959

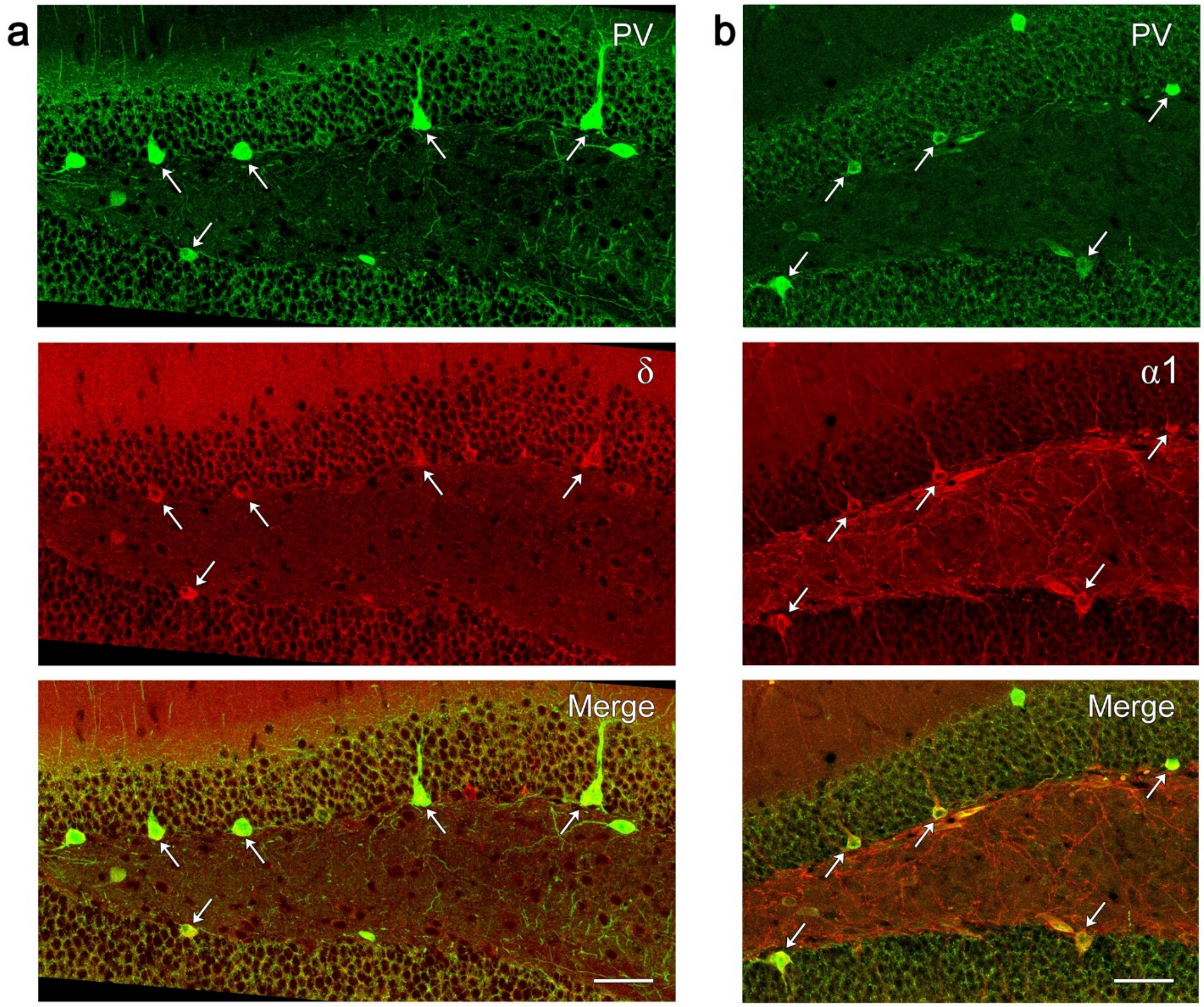


**Supplementary Figure S1** | **PV+ interneurons along the base of the granule cell layer express both the δ and α1 subunits. a,** The majority of PV+ interneurons along the inner border of the granule cell layer express the δ subunit (examples at arrows). **b,** Virtually all PV+INs along the granule cell border also express the α1 subunit (examples at arrows). Such labeling supports the localization of δ and α1 subunits in these PV+ interneurons and their possible subunit partnership. Scale bars: **a,b** = 50 μm.

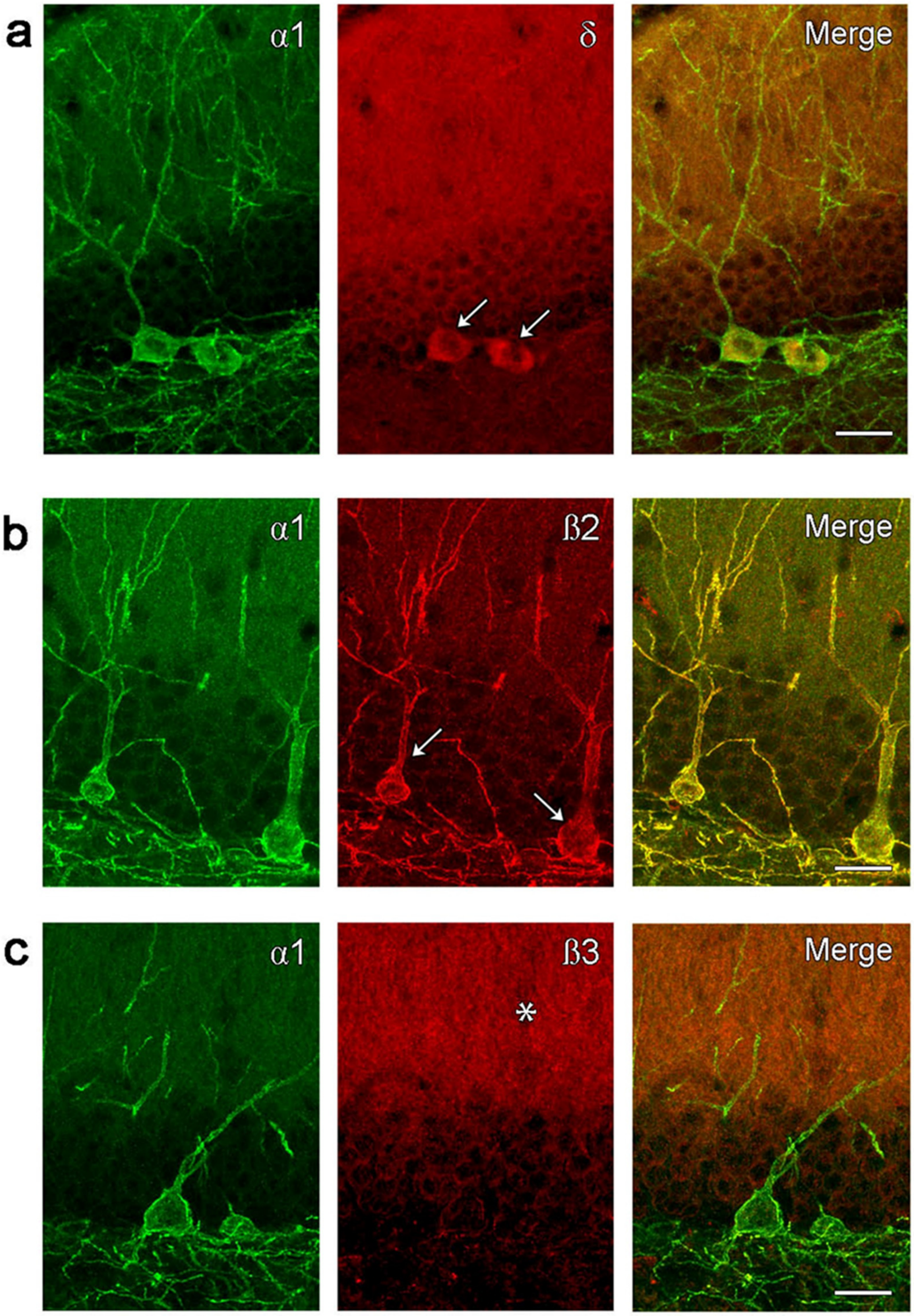


**Supplementary Figure S2 | α1 subunit-labeled interneurons at the base of the granule cell layer express δ and β2 subunits.** **a,** δ subunit labeling (arrows) is evident in the cell bodies of α1-labeled interneurons. **b,** β2 subunit labeling (arrows) is extensively co-localized on the surface of α1-labeled cell bodies and dendritic processes. **c,** β3 subunit shows little localization on the α1-labeled interneurons and is concentrated in small punctate structures in the dentate molecular layer (*). These localization patterns support a receptor subunit partnership of δ, α1 and β2 in these interneurons. Scale bars: **a,b,c** = 20 μm.

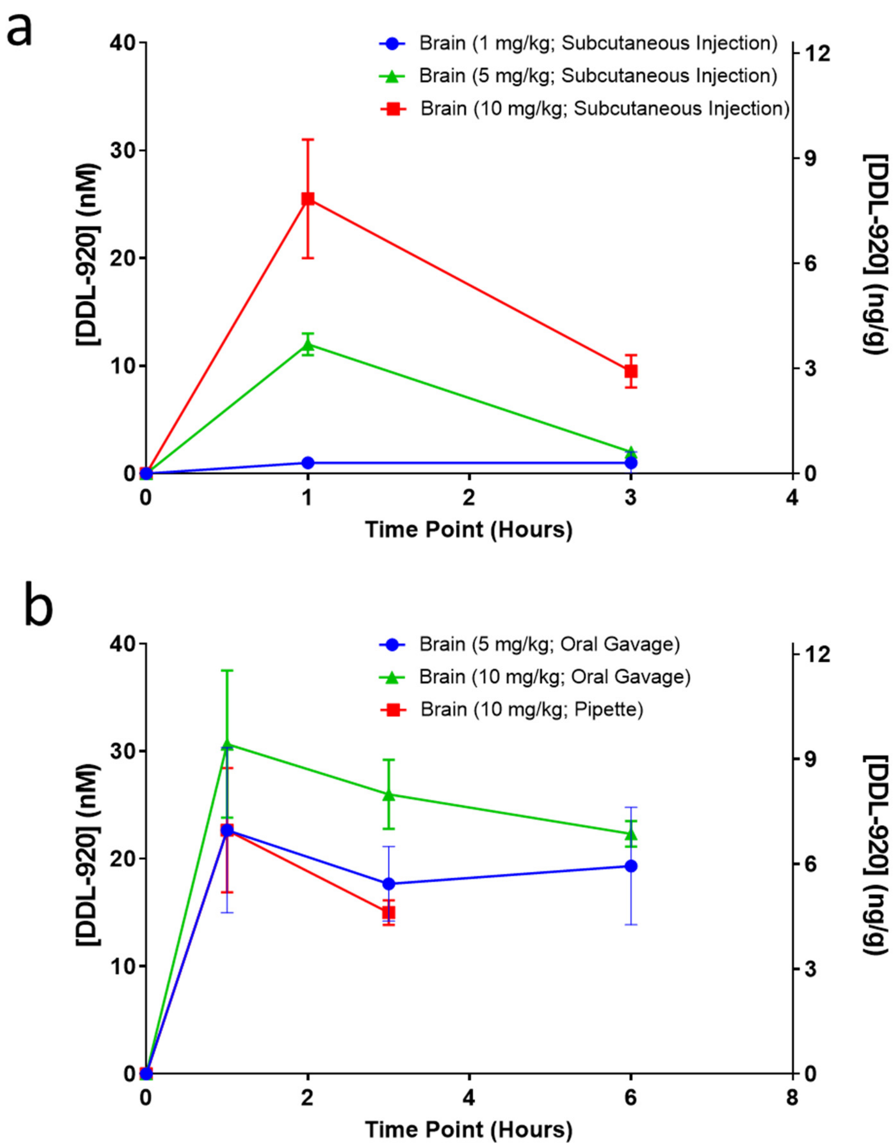


**Supplementary Figure S3 | DDL-920 PK studies by subcutaneous, oral gavage, or oral pipette administration. a,** Brain concentrations and levels of DDL-920 after subcutaneous administration at 1, 5 and 10mg/kg; **b,** Brain concentrations and levels of DDL-920 after oral gavage at 5 and 10 mg/kg, or by pipette administration at 10 mg/kg.

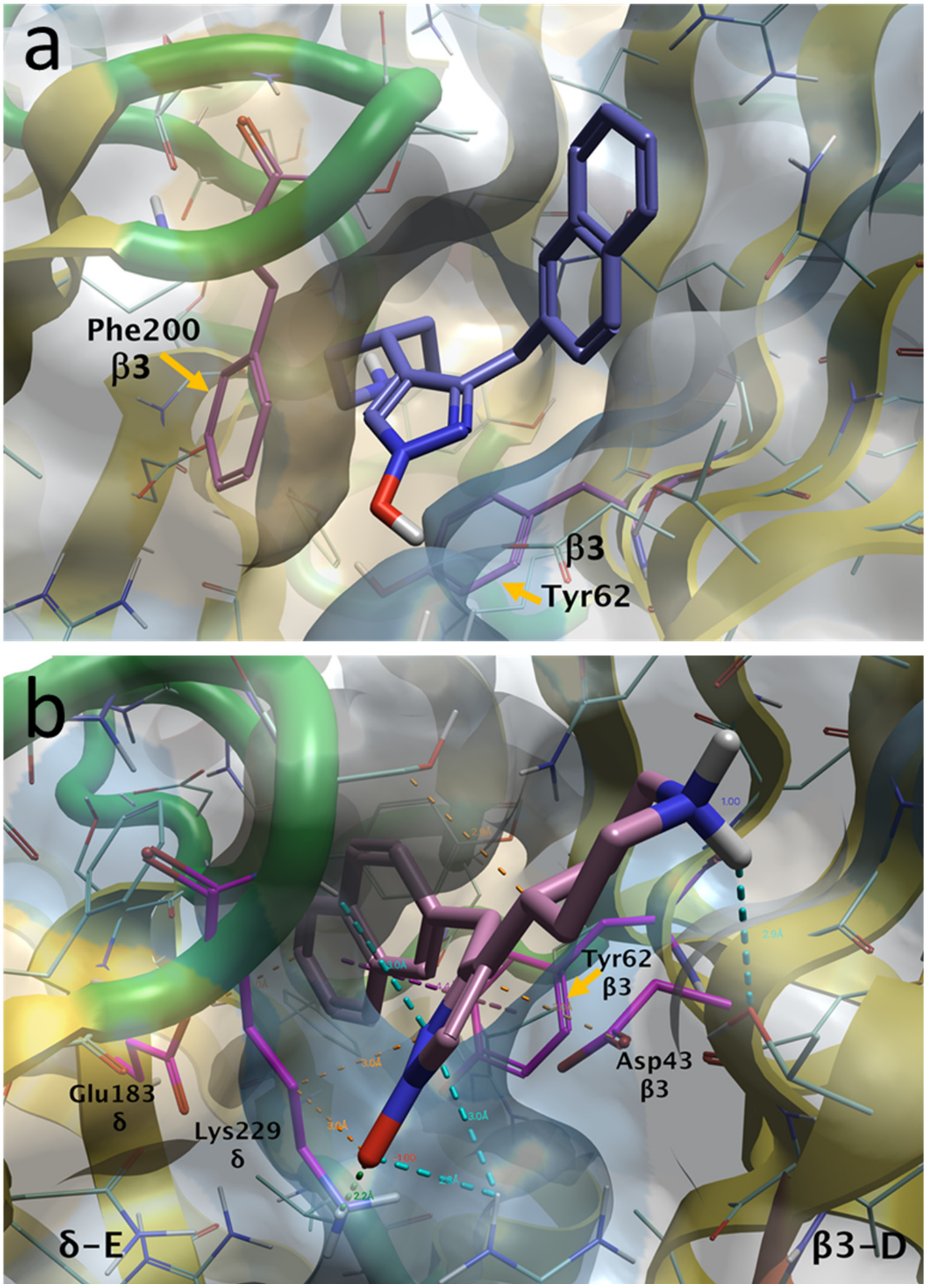


**Supplementary Figure S4 | DDL-920 docking to GABA$_A$Rs. a,** Docking of DDL-920 to the β3 homopentameric GABA$_A$R (pdb: 4COF) gave a good lead finder (LF) rank score (-10.34) and shows key interactions with Phe-200 and Tyr-62 of the b subunit; **b,** Docking to the β3δ heteropentameric GABA$_A$R structure (pdb: 7QND) gave a better LF rank score (-11.21) and shows how DDL-920 interacts with both the δ subunit (δ-E) and with the β subunit (β3-D). The key interaction of DDL-920 for β3-subunit was with Tyr-62 and Asp-43, while key interaction for the δ-subunit was with Lys-229 and Glu-183.

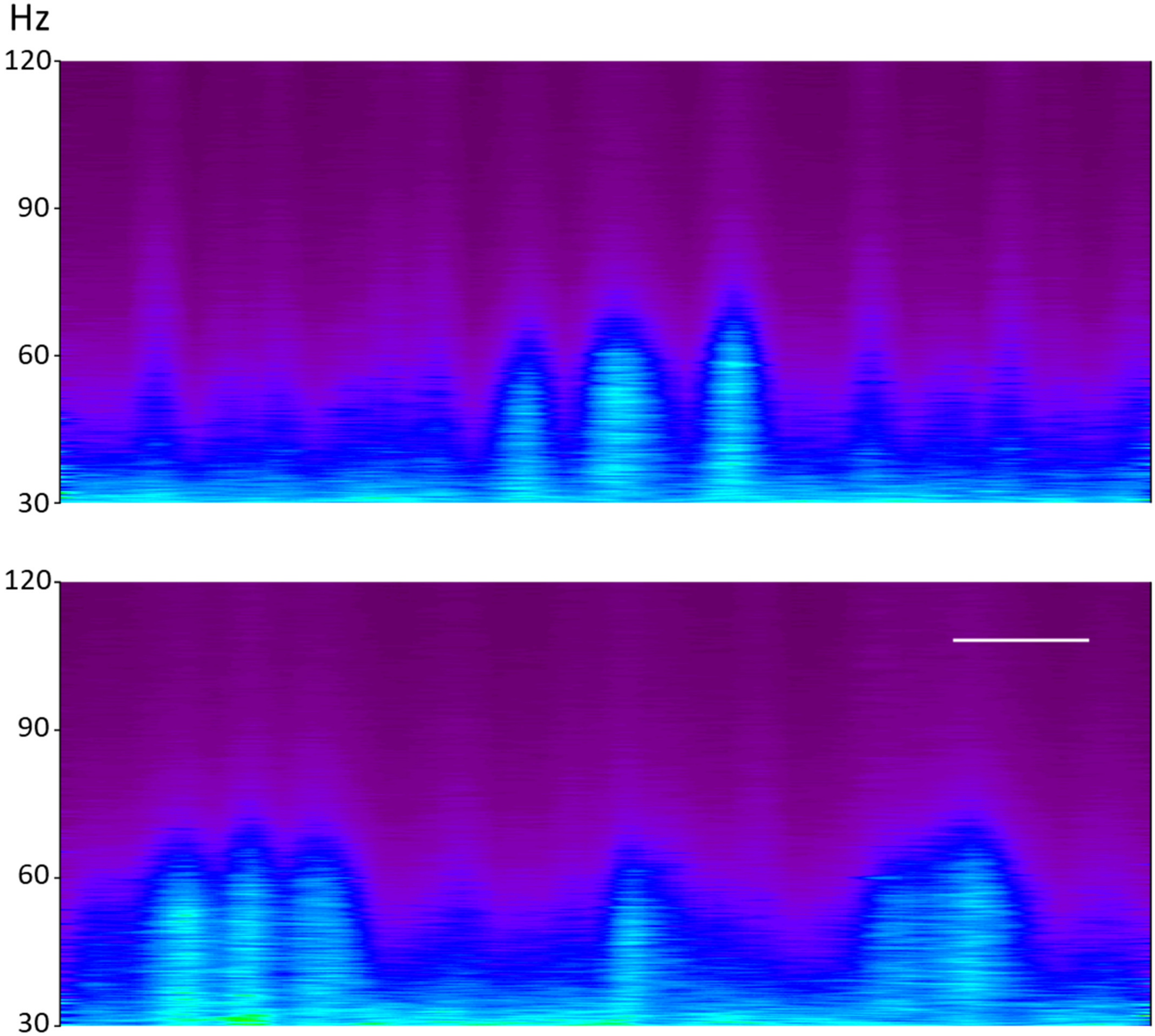


**Supplementary Figure S5 | Related to Figure 3 | Spectrograms of 4 hr recording periods of γ-oscillations.** *Top*: Spectrogram of the hippocampal γ-oscillations 4 hrs before a 10 mg/kg DDL-920 SQ injection on Day 0 (Figure 3) of a WT mouse. *Bottom*: Spectrogram of the hippocampal γ-oscillations for a duration 4 hrs commencing about 20 min after a 10 mg/kg DDL-920 SQ injection on Day 0 (Figure 3) of a WT mouse. Calibration bar is 30 min, and refers to both panels. For color scale values, please see Supplementary Figure 6.

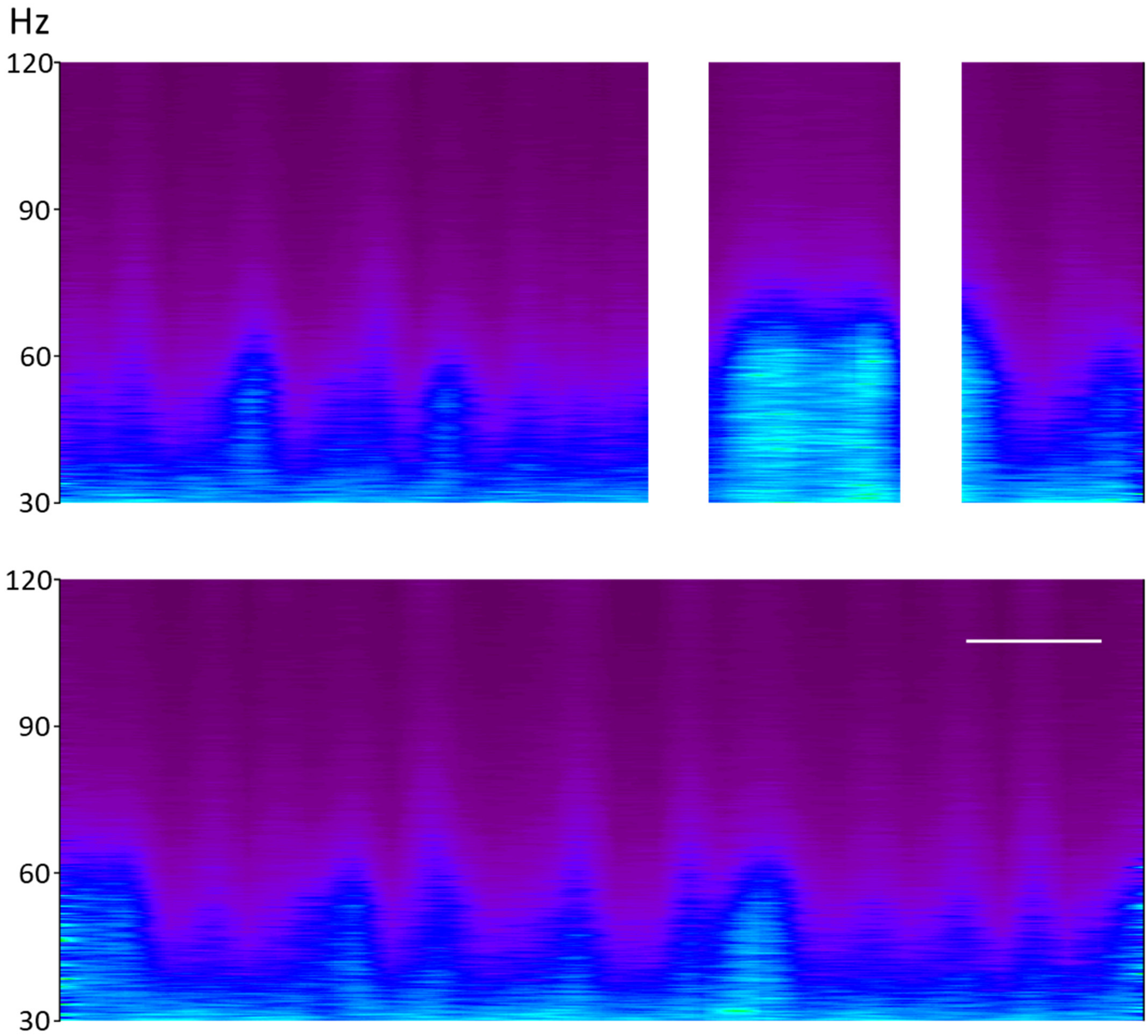


**Supplementary Figure S6 | Related to Figure 3 | Spectrograms of 4 hr recording periods of γ-oscillations.** *Top*: Spectrogram of the hippocampal γ-oscillations 4 hrs before a SQ saline injection on Day 3 (Figure 3) of a WT mouse. White bars indicate periods where the spectrograms were biased by excessive electrical/mechanical noise in the recording apparatus. *Bottom*: Spectrogram of the hippocampal γ-oscillations for a duration of 4 hrs commencing about 30 min after SQ saline injection on Day 3 (Figure 3) of a WT mouse. Calibration bar is 30 min, and refers to both panels. For color scale values, please see Supplementary Figure 6.

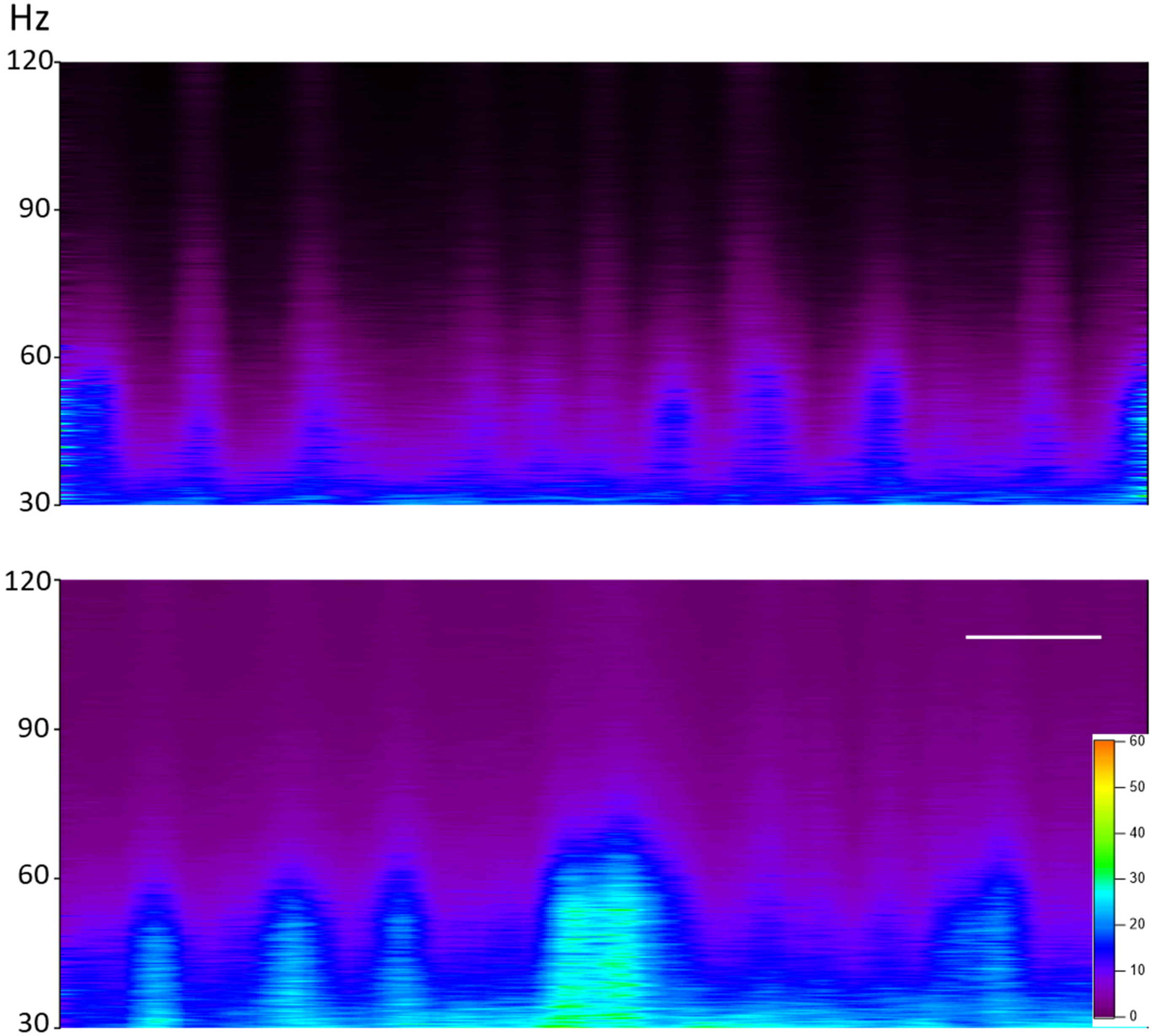


**Supplementary Figure S7 | Related to Figure 3 | Spectrograms of 4 hr recording periods of γ-oscillations.** *Top*: Spectrogram of the hippocampal γ-oscillations for 4 hrs before a 10 mg/kg DDL-920 SQ injection on Day 24 (Figure 3) of a WT mouse. *Bottom*: Spectrogram of the hippocampal γ-oscillations for a duration of 4 hrs commencing about 15 min after a 10 mg/kg DDL-920 SQ injection on Day 24 (Figure 3) of a WT mouse. Calibration bar is 30 min, and refers to both panels. Color scale values are in $\mu V^2$.

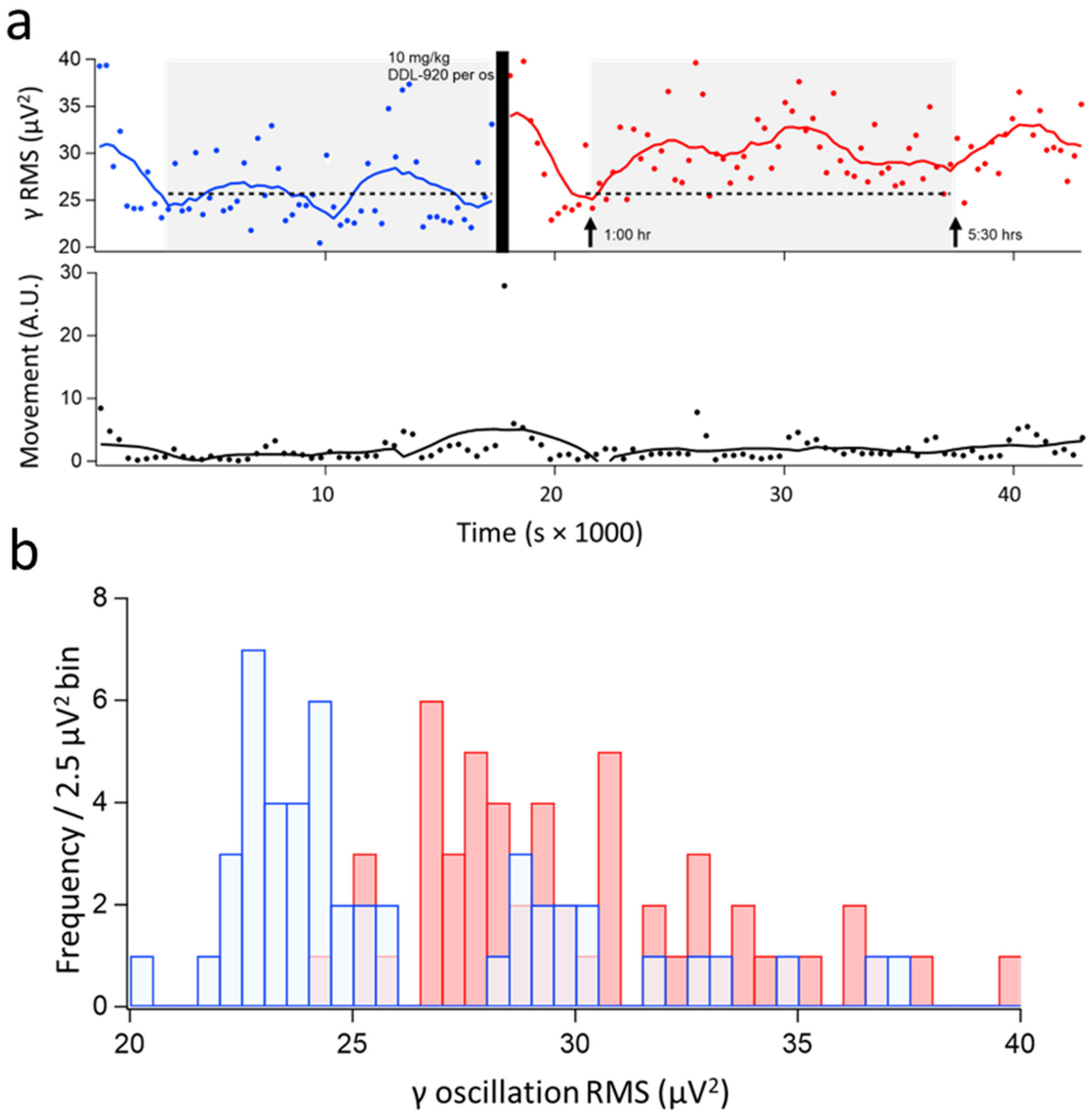


**Supplementary Figure S8** | **Related to Figure 4a-c. The power (RMS) of γ-oscillations in an AD mouse following oral pipette administration of DDL-920 (10 mg/kg). a,** *Top panel*: The average RMS of the γ-oscillations was calculated every 5 min for a period of 12 hrs (from 6:00 to 18:00 hrs). The drug administration was done at the time indicated by a black vertical bar. The dashed line before the DDL-920 administration is the linear fit to the ~4 hrs of recordings (*gray box*) except for the first 45 min of the recording when the animals are usually winding down from the time spent in the dark. The same fitted line was extended to the post drug administration period, allowing for 1 hr of recovery after the animal has been handled during the drug administration. It appears that the effect of DDL-920 on γ-oscillation power lasts more than 4.5 hrs, which constitutes a good estimate of its pharmacodynamics. *Bottom panel*: The corresponding motor activity of the mouse, also averaged every 5 min, as derived from the video recordings, is plotted on the same x-axis to illustrate no change in behavior. **b,** Histogram of the points during the shaded periods of **a**. The RMS means (± SD) were 26.1 ± 4.1 $\mu V^2$ prior to DDL-920 administration, and 30.1 ± 3.5 $\mu V^2$ after the drug administration ($p = 1.13e-6$, t-test).

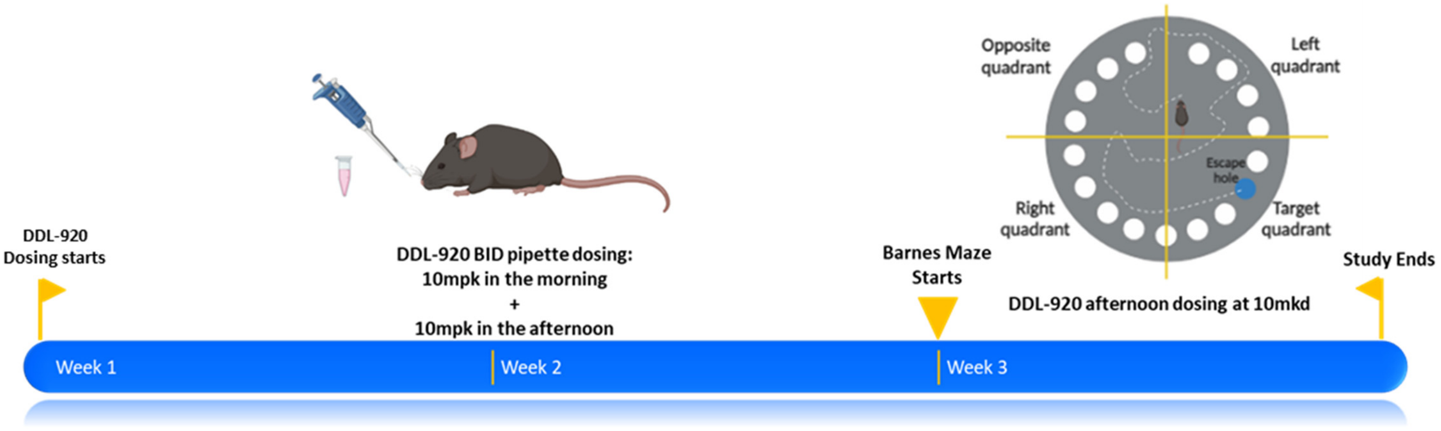


**Supplementary Figure S9 | Efficacy testing for DDL-920 on memory in the AD mouse model**. DDL-920 testing in the ApoE4-TR:5xFAD mice was done by oral pipette administration at 10 mg/kg twice daily (BID) for 2-weeks followed by Barnes Maze memory testing during the third week. The mouse, pipettor, vial, and Barnes Maze in the figure have been adapted from www.biorender.com.

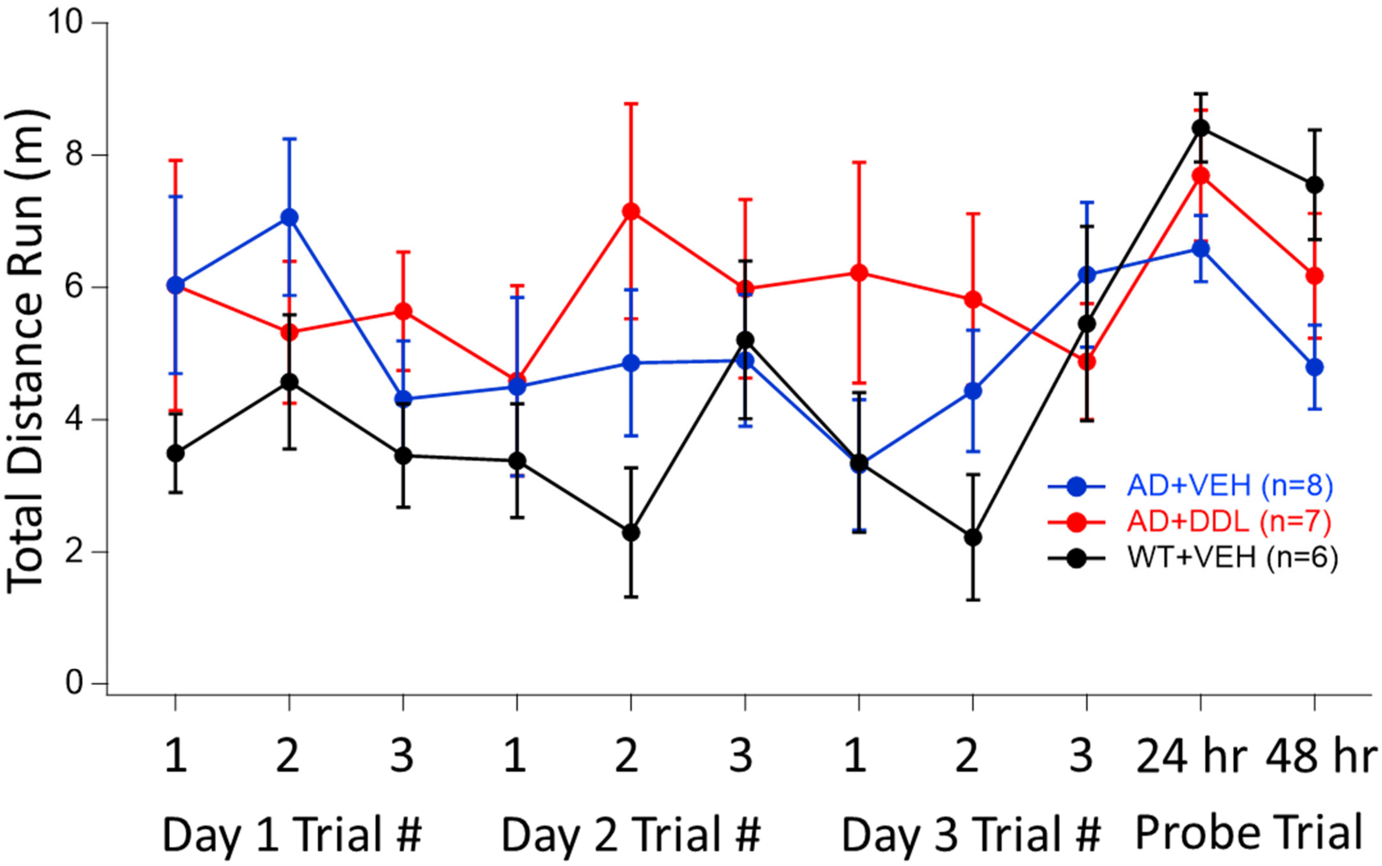


**Supplementary Figure S10 | Mouse motility in the Barnes maze during the training period and the probe trials**. The total distance run by the three groups of mice in the Barnes maze did not differ significantly between the groups, neither during the training trial periods, nor during the 24 hr or 48 hr probe trials. Points represent means, error bars are SEM.

| Property or ADME Parameter | Measured | Desired |
|---|---|---|
| Molecular weight [g/mol] | 342.86 | >160 & <480 |
| Ratio of sp3 hybridized C : total C count (Fraction Csp3) | 0.32 | >0.25 |
| Number of rotatable bonds | 3 | <8 |
| Number of H-bond acceptors | 3 | <7 |
| Number of H-bond donors | 2 | <3 |
| Kinetic solubility [μM] | >100 | > 5 |
| Microsomal stability [$t_{1/2}$ (min)] | >120 | >60 |
| Plasma stability [$t_{1/2}$ (min)] | >150 | >180 |
| Plasma binding [$F_{unbound}$ (%)] | 14.6 | >10 |
| BBB permeability PAMPA [$P_m$] | 6.0 | >0.85 |
| Brain: Plasma ratio | 0.714 | >0.5 |
| Brain tissue binding [$F_{unbound}$ (%)] | 34 | >10 |
| Oral (pipette) brain pharmacokinetics [$C_{max}$ (nM)] | 23 | |
| Unbound brain concentration [$C_{max}$ (nM)] | 8 | |
| In vitro efficacious dose [nM] | 1 | |

**Supplementary Table S1 | Physico-chemical properties[1] and in vitro absorption, distribution, metabolism and excretion (ADME) data[2-5] for DDL-920.** The drug-like properties of DDL-920 have desirable properties for further drug development.
